# Identification of motif-based interactions between SARS-CoV-2 protein domains and human peptide ligands pinpoint antiviral targets

**DOI:** 10.1101/2022.10.07.511324

**Authors:** Filip Mihalic, Caroline Benz, Eszter Kassa, Richard Lindqvist, Leandro Simonetti, Raviteja Inturi, Hanna Aronsson, Eva Andersson, Celestine N. Chi, Norman E. Davey, Anna K. Överby, Per Jemth, Ylva Ivarsson

## Abstract

The infection and replication cycle of all viruses depend on interactions between viral and host proteins. Each of these protein-protein interactions is therefore a potential drug target. These host-virus interactions often involve a disordered protein region on one side of the interface and a folded protein domain on the other. Here, we used proteomic peptide phage display (ProP-PD) to identify peptides from the intrinsically disordered regions of the human proteome that bind to folded protein domains encoded by the SARS-CoV-2 genome. Eleven folded domains of SARS-CoV-2 proteins were found to bind peptides from human proteins. Of 281 high/medium confidence peptides, 23 interactions involving eight SARS-CoV-2 protein domains were tested by fluorescence polarization, and binding was observed with affinities spanning the whole micromolar range. The key specificity determinants were established for six of these domains, two based on ProP-PD and four by alanine scanning SPOT arrays. Finally, two cell-penetrating peptides, targeting Nsp9 and Nsp16, respectively, were shown to function as inhibitors of viral replication. Our findings demonstrate how high-throughput peptide binding screens simultaneously provide information on potential host-virus interactions and identify ligands with antiviral properties.

## Introduction

The coronavirus disease 2019 (COVID-19) pandemic caused by the severe acute respiratory syndrome coronavirus 2 (SARS-CoV-2) virus has overwhelmed hospitals around the world and pushed public health facilities to their limits in the past years. Rapid vaccine development has drastically reduced this burden, but given the high rate of vaccine breakthrough cases, better post-infection therapeutic interventions are needed. Currently, many academic and industrial laboratories are working to develop new drugs or testing the effect of repurposing existing drugs against SARS-CoV-2. For example, molnupiravir is a nucleoside analogue targeting viral RNA-dependent RNA polymerase^1^, and paxlovid is a combination of two protease inhibitors, one previously used against human immunodeficiency virus (HIV) and the other developed to target the 3C-like protease of coronaviruses^2^. However, to be prepared for future epidemics and pandemics, it is important to continue developing new broad-spectrum antivirals such that emerging viral threats can be treated directly and immediately with approved drugs, preferably administered orally. This is particularly important for SARS-like coronaviruses, given the efficiency of human-to-human transmission, and has been demonstrated by recurrent outbreaks: severe acute respiratory syndrome coronavirus (SARS-CoV) in 2003, Middle East respiratory syndrome-related coronavirus (MERS-CoV) in 2012 and SARS-CoV-2 in 2019^3^.

While most drugs work by inhibiting enzymes, the pharmaceutical industry is turning its attention to the more challenging but largely untouched area of protein-protein interaction interfaces. To this end, several approaches have been developed to identify novel drug targets by mapping the virus-host protein-protein interactome^4–7^. In a particular class of protein-protein interactions, an intrinsically disordered region (IDR) of a protein interacts with a folded domain of the binding partner. The most common interaction modules in IDRs are short linear motifs (SLiMs). SLiMs are usually less than 10 amino acid residues long and degenerate, with only 3-4 residues accounting for most of the specificity and affinity. Consequently, SLiMs can evolve *ex nihilo* through accumulation of a few mutations. Because viruses evolve relatively rapidly, viral mimicry of host protein SLiMs has been proposed as a commonly used strategy, and several examples have been described across most viral clades^6–8^. Proteins expressed from viral genomes therefore often contain SLiMs which compete with native host protein-protein interactions to hijacking cellular machinery. Conversely, folded domains of the viral proteome can interact with both human and viral SLiMs^8–11^. Each scenario provides a drug target, *i.e*., the folded domain of human or viral protein can be targeted with a peptide or peptidomimetic.

The SARS-CoV-2 proteome consists of 29 proteins, encoded by 14 open reading frames (ORF)^5^. The first two reading frames, ORF1a and ORF1b, encode 16 non-structural proteins (Nsp1 to Nsp16), formed after post-translational proteolytic cleavage by viral proteases. In addition, the SARS-CoV-2 genome codes four structural proteins: spike (S), nucleocapsid (N), membrane (M), and envelope (E), as well as nine accessory proteins (ORFs 3a, 3b, 6, 7a, 7b, 8, 9b, 9c, and 10) **(Figure 1)**. Non-structural proteins facilitate viral mRNA replication and translation. Nsp1 is responsible for suppressing host translation while simultaneously promoting viral mRNA translation^12^. Nsp2 promotes viral evasion of the innate immune response^13^. Nsp3 serves as a central hub coordinating different steps of viral replication^14,15^. It consists of several stably folded globular domains, namely two ubiquitin-like domains (Ubl1, Ubl2), an ADP-ribose-1’’-(phosphatase) hydrolase (ADRP), three SARS-unique domains (SUD-N, SUD-M and SUD-C), a papain-like protease domain (PLpro) and a nucleic acid binding domain (NAB). In addition to Nsp3, Nsp4 and Nsp6 are also involved in double membrane vesicle formation and organization^16,17^. Nsp5 serves as the main protease (MPro), that proteolytically processes the ORF1a and ORF1b into final proteins^18^. Nsp7, Nsp8, and Nsp12 form the core replication complex with Nsp12 being the RNA dependent RNA polymerase (RdRp) and Nsp7/Nsp8 forming a hexadecameric complex, that enhances processivity^19–21^. In addition, Nsp9 also associates with the replication complex to promote 5’- capping of viral RNA^22^. The addition of a 5’ cap to the viral RNA is crucial for RNA stability as well as efficient translation and requires several steps, with the Nsp10/Nsp14 and Nsp10/Nsp16 complexes also contributing to the final steps of the process^22^. In both cases, Nsp10 activates the catalytic activity of Nsp14 and Nsp16. Apart from its function in the 5’- capping process, Nsp14 also functions as an exoribonuclease facilitating the proofreading function of the replication complex. Finally, Nsp13 is a helicase, that unwinds the RNA and promotes efficient transcription^23^, while Nsp15 serves as a uridine-specific endoribonuclease that facilitates the evasion of the host immune response^24^.

**Figure 1:**
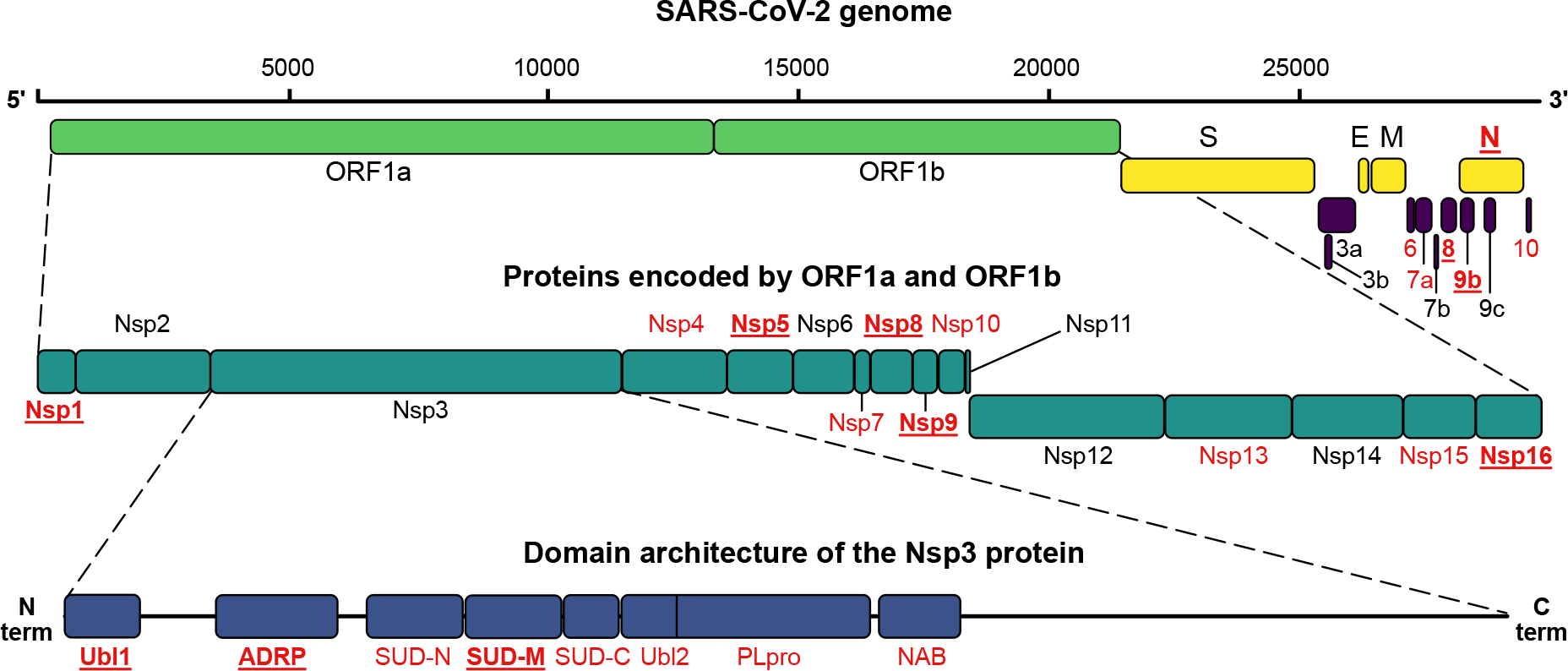
Organization of the SARS-CoV-2 genome and proteome. Proteins and protein domains that were successfully expressed in this study are shown in red. Proteins and protein domains that enriched ligands in ProP-PD selections are shown in bold and underlined. The figure is adapted from Gordon et. al.^5^

To investigate the virus-human protein-protein interactome of folded protein domains encoded by the SARS-CoV-2 genome and SLiMs in the human proteome, we created a collection of protein domain expression constructs from SARS-CoV-2 proteins. These protein domains were used as baits in proteomic peptide phage display (ProP-PD) screens against a Human Disorderome (HD2) peptide library that displays 1 million peptides from the human proteome^25^. The screen identified peptide-binding SARS-CoV-2 proteins and human peptide ligands that were validated with binding assays and tested in a SARS-CoV-2 infection assay. Our results suggest that protein domains of SARS-CoV-2 may serve as targets for the design and development of peptidomimetic antivirals.

## Results

### Large-scale identification of SLiM-based interactions and interaction motifs of SARS-CoV-2 proteins

We generated a collection of 31 expression constructs of known or predicted modular domains from 22 SARS-CoV-2 proteins, including two catalytically inactivated protease variants (**Figure 1; Table S1**). The domains were expressed as GST-tagged fusion proteins, and 26 of the 31 protein constructs were produced in sufficient quantities to be used as baits in phage display selections. The GST-fusion proteins were immobilized for triplicate selections in a 96-well plate and used in ProP-PD selections using our HD2 peptide phage-display library. This library displays 16 amino acid overlapping peptides that tile the intrinsically disordered regions of the human proteome on the surface of M13 phage^25^. The peptide-coding regions of the binding-enriched phage pools were analyzed by next-generation sequencing (NGS). Using previously established quality metrics (peptides found in replicate selections, overlapping peptides identified, high NGS score and/or containing motifs) we found 281 high/medium (8/273) confidence peptides from 239 proteins interacting with 11 SARS-CoV-2 domains of nine viral proteins (Nsp1, Nsp3 (Ubl1, ADRP and SUD-M), Nsp5, Nsp8, Nsp9, Nsp16, Orf8, Orf9b and N NTD) (**Figure 1, Table S2**). The vast majority (118) of identified interactions involve Nsp9, followed by Nsp1 (47) and the catalytically inactivated Nsp5 Mpro (32). Based on the peptides identified by ProP-PD selections, consensus motifs could be established for two proteins (Nsp5 [FLM][HQ][AS] and Nsp9 G[FL]xL[GDP]; **Figure 2**). For Nsp5 (Mpro), the ligands may serve as potential substrates because the recognition motif resembles the preferred proteolytic site of the protease (LQ↓[GAS])^26^.

**Figure 2:**
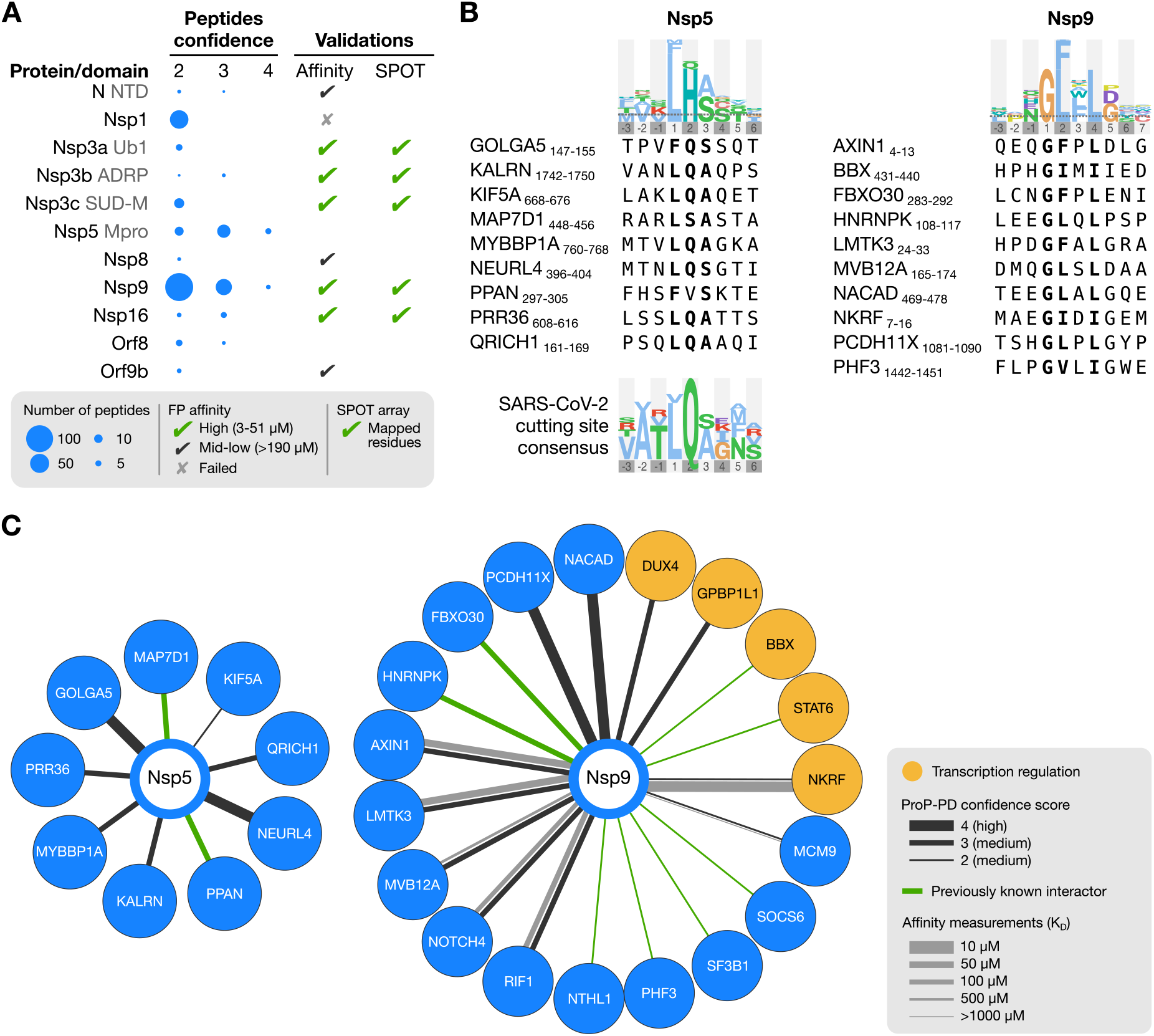
Outline of the results. **(A)** Overview of the results for the 11 SARS-CoV-2 protein domains that enriched peptides with medium (2, 3) and high (4) confidence scores in ProP-PD selections. The number of peptides is proportional to the area of the blue circles. Overall validation of affinity with fluorescence polarization and SPOT array is indicated. **(B)** The motifs obtained from the selections with the SARS-CoV-2 proteins Nsp5 and Nsp9 are highlighted in bold in the alignments of the representative peptides. The consensus motif for the Nsp5 (Mpro) protease cleavage site is shown below the alignment^26^. **(C)** Subset of identified interactions of SARS-CoV-2 Nsp5 and Nsp9. The thickness of dark lines shows ProP-PD confidence scores for the interactions, and the thickness of grey lines the *K*_D_ values obtained from the fluorescence polarization assays. Human proteins that contain the enriched GO terms associated with the transcriptional regulation (GO:0003700, GO:0000981, GO:0000976, GO:0001228, GO:0000978, GO:0000977, GO:0006355) are highlighted in orange.

The enrichment analysis of the Gene Ontology (GO) terms for the combined dataset revealed enrichment of ligands associated with biological processes related to transcriptional regulation (**Table S3;** q < 0.001; PepTools). By comparing with the information available in protein interaction databases (collected August 2022, See Materials and Methods for details) we found that eighteen of the protein-protein interactions identified here are supported by reported interactions from previous studies (**Figure 2, Table S2**). The low overlap with other host-SARS-CoV-2 protein-protein interaction studies likely reflects technical differences between the approaches^27^. Here, we specifically searched for SLiM-based interactions, which are often missed by other methods^25^.

We selected 27 interactions for validation in a fluorescence polarization (FP) based binding assay, in which we first establish a valid fluorescein (FITC) labelled probe peptide, which was subsequently displaced in a second experiment with unlabelled peptide(s). The following SARS-CoV-2 protein domains were included in these binding experiments: Nsp1, Nsp3 Ubl1, Nsp3 ADRP, Nsp3 SUD-M, Nsp3 NAB, Nsp8, Nsp9, Nsp16, Orf9b, and N-NTD (**Table S4**). For Nsp1 and Nsp3 NAB we did not detect binding with the probe peptides tested whereas weak, non-saturating binding was observed for Nsp8, Orf9b and N NTD, indicating that the tested interactions are of low affinity (**Figure S1, Table S4**). In contrast, we conclusively confirmed the peptide binding capacity of five SARS-CoV-2 domains: Nsp3 Ubl1, Nsp3 ADRP, Nsp3 SUD-M, Nsp9 and Nsp16, as described below.

### Characterization of SLiMs binding to distinct domains of Nsp3

Coronavirus replication occurs primarily in double-membrane vesicles, derived from the host endoplasmic reticulum membrane, that provide protection from the host immune response^28^. Recently, the hexameric Nsp3 assembly was shown to form pores that span the double membrane and serve as a vital connection between the site of viral RNA synthesis inside the vesicle and the site of viral RNA translation in the cytoplasm^29^. Despite great recent efforts to map and characterize the SARS-CoV-2 interactome, the Nsp3 has been largely neglected due to its size and complexity, and therefore the functions of the various domains are poorly understood^5,30^. We expressed and purified all globular domains of Nsp3 and subjected them to Pro-PD selections **(Figure 1)**. Phage selections yielded a set of medium confidence ligands for Nsp3 Ubl1 (6 peptides), ADRP (3 peptides) and Nsp3 SUD-M (14 peptides) but without a clear consensus motif for any of the domains, in part due to the small number of retrieved peptides **(Table S2)**. We determined the equilibrium dissociation constant (*K*_D_) using the FP assay for at least two human peptides for each of the three globular Nsp3 domains **(Figure 3A, Figure S1)**, which confirmed the binding of six of the tested peptides with affinities in the range of 20 to 300 μM (**Table S4**).

**Figure 3:**
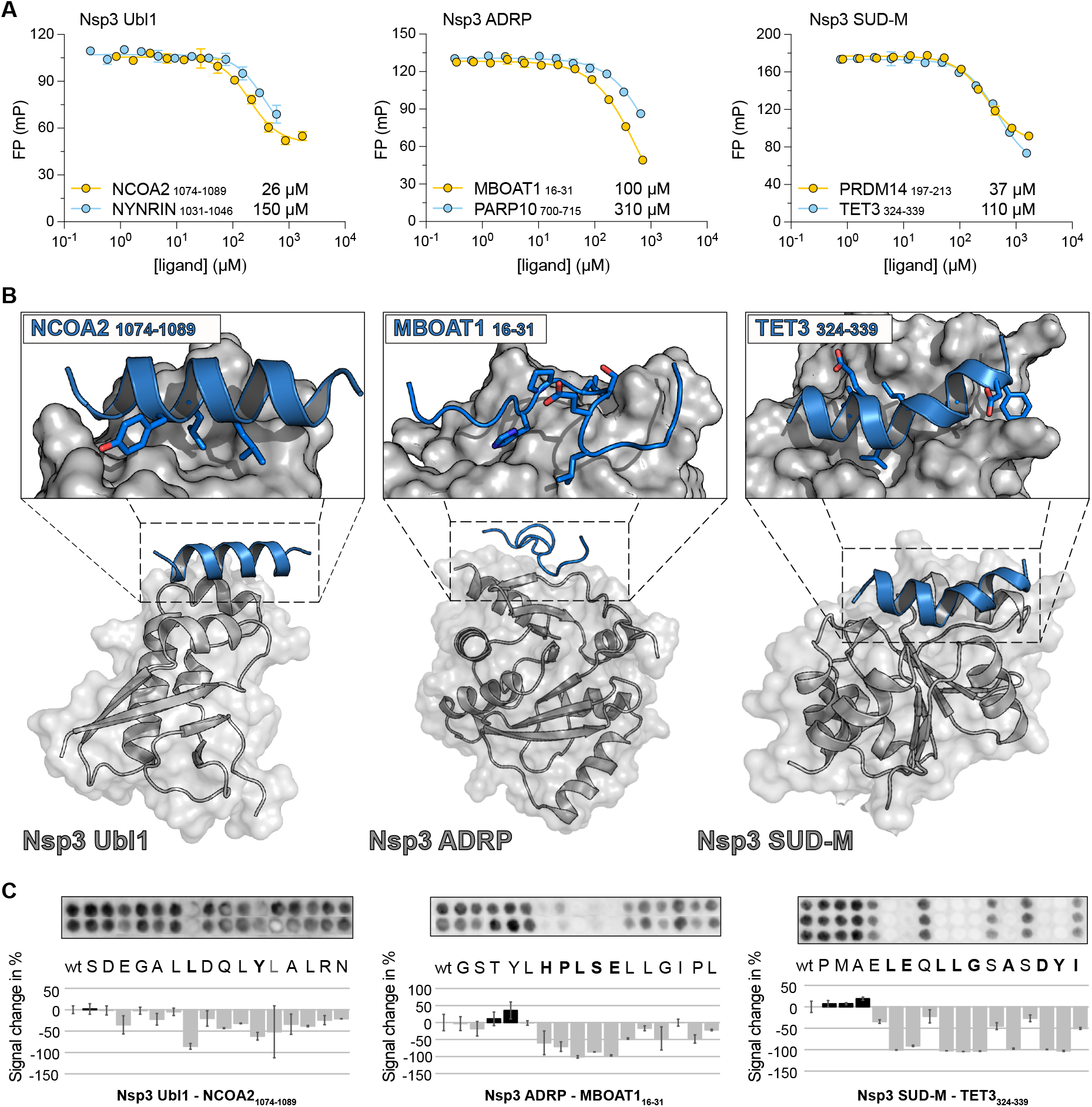
Biophysical analysis of the interactions between Nsp3 Ubl1, Nsp3 ADRP, and Nsp3 SUD-M, with peptide ligands from human proteins. **(A)** Fluorescence polarization-monitored displacement experiments measuring the affinity between globular domains of Nsp3 and peptide ligands from disordered regions of the human proteome identified by phage display. The *K*_D_ values are shown next to each peptide and in **Table S4**. **(B)** ColabFold structural predictions for the interaction between the globular domains of Nsp3 and the peptide ligands. The globular domains of Nsp3 are shown in grey, whereas the peptides are shown as blue ribbons. The ColabFold pLDDT confidence scores were high for the globular domains of Nsp3 (>90) but varied widely for peptide predictions (30-80) and are shown in **Figure S3**. **(C)** SPOT array alanine scans for the indicated peptides. Residues involved in binding are shown in bold. Signal intensities were normalized to wild type (wt) and presented as average percent signal change. Error bars indicate one standard deviation from the average.

Three peptides derived from the protein NYNRIN (NYNRIN_1031-1046_; CPSLSEEILRCLSLHD), the Nuclear receptor coactivator 2 (NCOA2) (NCOA2_1074-1089_; SDEGALLDQLYLALRN), and from the Chromatin complexes subunit BAP18 (BAP18_40-55_; AKWTETEIEMLRAAVK) were selected for affinity measurements with the Nsp3 Ubl1. The measurements confirmed the binding of Nsp3 Ubl1 to NYNRIN_1031-1046_ and NCOA2_1074-1089_ (**Figure 3A, Table S4**), but not to BAP18_40-55_. The two binding peptides shared some sequence similarity, and possessed a putative LxxLxL motif, which is known to adopt alpha helical conformation upon binding^31,32^. One such interaction exists between the NCBD domain of CBP/p300 and the CID domains of the NCOA2 protein family^33^. We therefore tested the binding of Nsp3 Ubl1 to the entire CBP interaction domain (residues 1071-1110) of NCOA2 as well as to the two human paralogs NCOA1 and NCOA3 (residues 924-965 and 1045-1086 respectively). We did not detect any displacement for the two paralogues suggesting that the conserved LxxLxL motif is not the main driver of the interaction **(Figure S1, Table S4)**. To clarify which residues of the NCOA2 peptide are involved in the interaction, we performed a peptide SPOT array alanine scan, which revealed that the leucine residues in position 1 (P1), together with a tyrosine in position P5, was critical for binding (**Figure 3C)**, indicating that the Nsp3 Ubl1 binding motif in the NCOA2_1074-1089_ is LxxxY. This finding clarified the lack of binding of NCOA1 and NCOA2 paralogues in which valines are located at the corresponding tyrosine position. Similarly spaced hydrophobic residues are also found in the other Nsp3 Ubl1 binding peptides (**Figure S2**).

To test how the peptides might bind to Nsp3 Ubl1 we performed an *in silico* prediction using ColabFold, which is based on AlphaFold2^34^. Both the NCOA2_1074-1089_ and the NYNRIN_1031-1046_ peptides were predicted with a high per-residue confidence score (pLDDT > 80) to form an alpha helix bound in the hydrophobic binding pocket between alpha helix 1 (α1) and helix 3 (α3) of Nsp3 Ubl1 (**Figure 3B; Figure S3**). Intriguingly, a recently reported high-affinity interaction between the disordered region of the SARS-CoV-2 nucleoprotein (N; 219-LALLLL-224) and Nsp3 Ubl1 also engages the same hydrophobic binding pocket of Nsp3 Ubl1^9^. In this case, the interaction is enhanced by an additional polar motif of N, that binds to Nsp3 Ubl1 in a distinct site. Therefore, to investigate if the ligands identified in this study directly compete with N for binding to Nsp3a Ubl1, we attempted to displace the FITC-labelled NCOA2_1074-1089_ peptide probe from the Nsp3 Ubl1-probe complex with full length N. However, the FP signal increased upon addition of N to the premixed Nsp3 Ubl1-probe sample as well as upon addition of N to the sample containing only the probe peptide, rendering the displacement experiment inconclusive (**Figure S1)**.

Next, we focused on the binding of peptides to the other two Nsp3 domains, ADRP and SUD-M. The peptides that bind to Nsp3 ADRP were derived from Lysophospholipid acyltransferase 1 (MBOAT1_16-31_; GSTYLHPLSELLGIPL) and Protein mono-ADP-ribosyltransferase PARP10 (PARP10_700-715_; DGGTDGKAQLVVHSA), with the MBOAT1 peptide being the ligand with higher affinity **(Figure 3A, Table S2** and **Table S4)**. The peptide SPOT array showed a strong signal for binding of the MBOAT1_16-31_ peptide to Nsp3 ADRP, suggesting that the central _21_-HPLSE-_25_ residues are critical for binding (**Figure 3C**, residues in bold). A similar _4_-YLSE-_7_ segment was also identified in AZIN2, an additional Nsp3 ADRP ligand found in the ProP-PD selection (**Figure S2**). The ColabFold predictions of the Nsp3 ADRP binding peptides did not converge with high confidence (pLDDT of ~30; **Figure S3**). However, manual inspection indicated that the peptide binding was restricted to the surface between the N-terminal beta sheet (β1) and the C-terminal alfa helix (α6) **(Figure 3B, Figure S3)**. Finally, we note that PARP10 is upregulated upon SARS-CoV-2 infection^35^ and is known to mono-ADP-ribosylate amino acid residues as a part of host response to viral infections^36^. The Nsp3 SUD-N (also called Mac1) domain of SARS-CoV-2 is in turn a mono-ADP-ribosylhydrolase^37^. Thus, although the affinity of the Nsp3 ADRP-PARP10 interaction is relatively low (*K*_D_ ≈ 300 μM), the interaction might be of relevance in the context of the virus counteracting the host’s response to viral infection.

The peptides used to study binding to Nsp3 SUD-M were from the PR domain zinc finger protein 14 (PRDM14_197-213_; FTEEDLHFVLYGVTPS) and Methylcytosine dioxygenase TET3 (TET3_324-339_; PMAELEQLLGSASDYI). These were the most enriched peptides in the selection against Nsp3 SUD-M, and of these two, PRDM14_197-213_ was the ligand with higher affinity (**Figure 3A**, **Table S2 and Table S4**). The ColabFold analysis of Nsp3 SUD-M in complex with either PRDM14_197-213_ or TET3_324-339_ exhibited moderate pLDDT scores (between 50 and 70, **Figure S3**). While the two peptides did not share a specific binding motif, the SPOT array alanine scan showed distinct patterns of alternating amino acid residues interacting with Nsp3 SUD-M for both peptides (**Figure 3C, Figure S4)**. In the case of Nsp3 SUD-M - TET3_324-339_ the ColabFold predictions and SPOT arrays converged allowing us to map a plausible binding interface, where TET3_324-339_ forms an alpha helix exposing the underlined motif _324_-PMAE**<u>LE</u>**Q**<u>LLG</u>**S**<u>A</u>**S**<u>DY</u>**I-_339_ for interaction with a hydrophobic pocket of SUD-M (**Figure 3B and 3C**).

Taken together we demonstrate that the Ubl1, ADRP, and SUD-M domains of Nsp3 possess the capacity to interact with SLiMs. The identified peptides bound with intermediate micromolar affinity and the binding sites on the peptides and the proteins were defined by alanine scans and docking predictions.

### Nsp9 binds to peptide ligands with consensus motifs, leading to the rearrangement of hydrophobic core and the disruption of dimerization

Nsp9 from SARS-CoV-2 is a 113 amino acid residue long RNA-binding protein with a stable fold that shares 98% sequence identity with its SARS-CoV homolog^38,39^. Recent reports have established that Nsp9 is an essential component of the replication and transcription complex where it interacts with the Nidovirus RdRp-Associated Nucleotidyltransferase (NiRAN) domain of the RNA-dependent RNA polymerase Nsp12^40^. In this model, the monomeric Nsp9 binds to NiRAN via a C-terminal alpha helix and facilitates the addition of the GpppA-RNA cap to the 5’-end of the newly synthesized mRNA^22^. The same helix has been suggested to facilitate the formation of the Nsp9 homodimer with the crucial _100_-GxxxG-_104_ motif at the dimer interface^41–45^.

In our ProP-PD selections, we identified 147 human Nsp9 binding peptides in which a G[FL]xL[GDP] motif consensus was enriched **(Figure 2; Table S2)**. We confirmed binding and determined the affinities for eight of the ligands, with the *K*_D_ values in the mid-micromolar range (**Figure 4A-B, Figure S1, Table S4**). The ligands with the highest affinity were peptides derived from the NF-kappa-B-repressing factor (NKRF_8-23_; AEGIDIGEMPSYDLVL), the protein kinase LMTK3 (LMTK3_22-36_; PAHPDGFALGRAPLA), and the Axin-1 (AXIN1_1-15_; MNIQEQGFPLDLGAS), which bound to Nsp9 with *K*_D_ values of 30-50 μM (**Table S4)**. Of these, NKRF has been found to interact with other SARS-CoV-2 proteins including Nsp1^46^ and Nsp10^47^ with the latter interaction thought to regulate interleukin-8 production. A peptide from the Neurogenic locus notch homolog protein 4 (NOTCH4_1605-1620_) displayed lower affinity, and bound to Nsp9 with a *K*_D_ ~130 μM. We further tested binding to the C-terminal peptide of Nsp9 (Nsp9_4237-4251_; LNRGMVLGSLAATVR), as it shares the GxxxG motif, but did not detect any measurable binding within the concentration range used (**Figure 4A**). To further dissect which residues of the peptides are essential for binding we performed a SPOT array alanine scan of the NKRF_8-23_ and LMTK3_22-36_ peptides. For the NKRF_8-23_ peptide the analysis confirmed the consensus motif, with a substantial decrease in binding intensity when either of the two glycines at the positions P1 and P5 of the motif or the isoleucine at the position P4 were mutated to alanine. A minor effect was also observed upon mutation of the isoleucine at the position P2 (**Figure 4D**). Based on these results, we adjust the motif to GΦxΦ[GDP], where Φ is a hydrophobic residue. To define the binding pocket of the motif, we performed ColabFold prediction for peptide binding to either monomeric or dimeric Nsp9. This analysis however, showed low confidence scores (pLDDT < 30; **Figure S5**) and did not converge on a similar binding mode for the three peptides.

**Figure 4:**
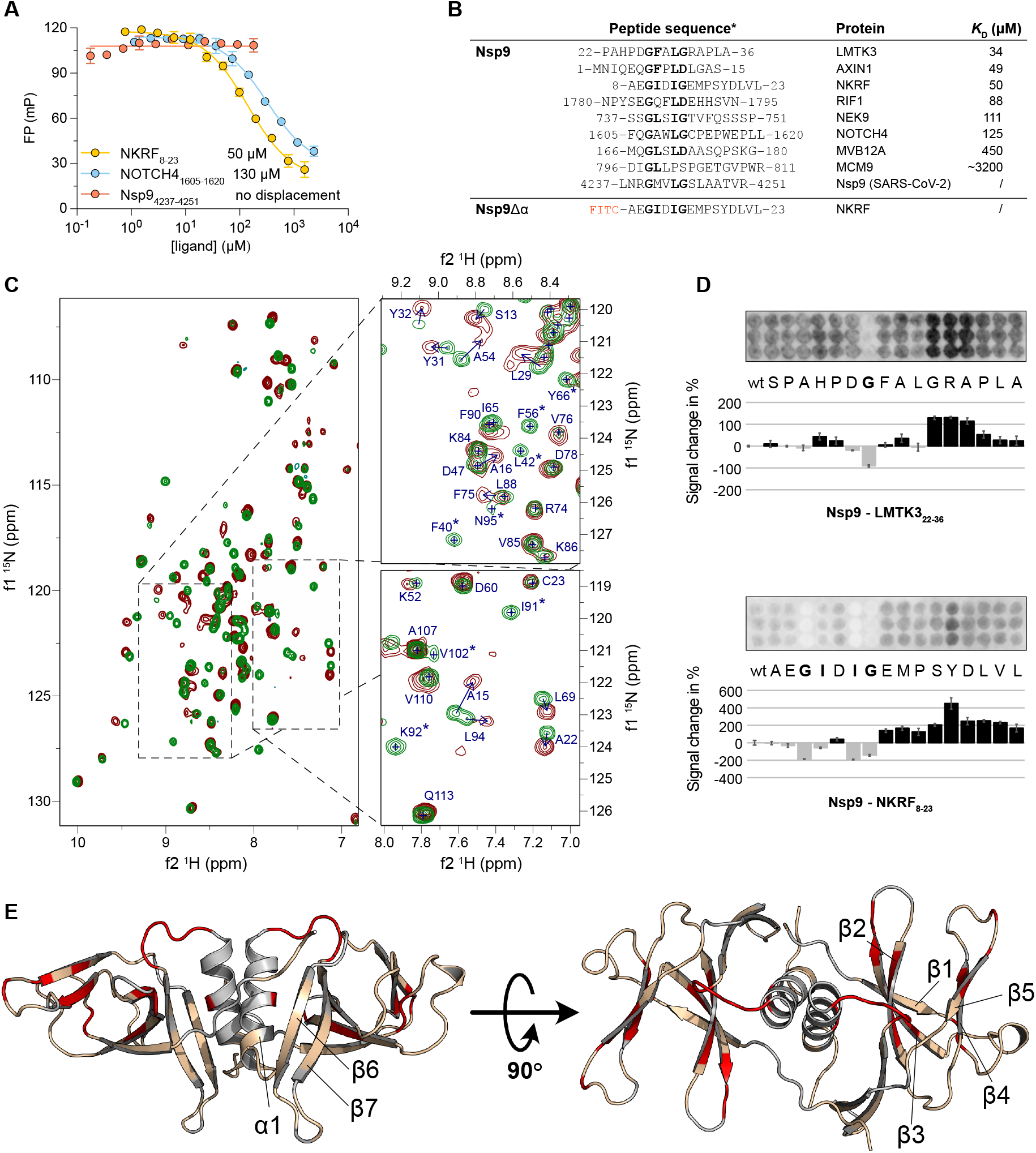
Interactions between Nsp9 and human peptide ligands are mediated by a GΦxΦ[GDP] motif and lead to rearrangement of the hydrophobic core of Nsp9. **(A)** Representative FP-monitored displacement experiments measuring the affinity between Nsp9 and three human peptide ligands. Affinities are shown next to each peptide. **(B)** Alignment of peptides identified by ProP-PD for which the affinities were measured. Residues, corresponding to the identified G&Φx&Φ[GDP] motif are shown in bold. For Nsp9Δα we did not observe saturation with the FITC-NKRF_8-23_ peptide. **(C)** HSQC experiments showing the ^1^H-^15^N peak shifts. The left panel shows the superposition of Nsp9 protein (green) and Nsp9 mixed with NKRF peptide (wine red). The right panels show representative examples of the observed chemical peak shift perturbations. The arrows indicate the directions of the chemical shift change and the asterisk indicates that the peaks disappeared after addition of the peptide, indicating a large perturbation of the chemical environment. The full spectra are shown in **Figure S6**. **(D)** SPOT array alanine scans for the indicated peptides. Residues involved in binding are shown in bold. Signal intensities were normalized to wild type (wt) and presented as percentage signal change. **(E)** Representation of the residues which displayed large change in chemical shift after addition of the NKRF_8-23_ peptide, as observed by NMR experiments. Residues that changed more than one standard deviation above the average are colored in red, the residues below this threshold are beige, and the residues whose peak shifts could not be unambiguously assigned are gray. The PDBid: 6wxd model of Nsp9 dimer was used for visualization.

Because the peptide ligands have a GΦxΦ[GDP] motif, that is also found in the C-terminal helix of Nsp9 and facilitates dimerization, we hypothesized that Nsp9-binding peptides might interact with the dimerization interface and therefore interfere with the dimer formation^41,48^. To address this question and map the residues involved in the interaction between Nsp9 and the peptides, we recorded nuclear magnetic resonance (NMR) ^1^H^15^N heteronuclear single quantum coherence (HSQC) spectra of ^15^N labeled Nsp9 at increasing concentrations of NKRF_8-23_, LMTK3_22-36_, and NOTCH4_1605-1620_, respectively. Well-resolved spectra, which were in excellent agreement with the previously described NMR assignments^44,48^, confirmed a folded protein and allowed us to monitor the changes in chemical shift perturbations upon addition of peptides **(Figure 4C, Figure S6, Table S5)**. The substantial overlap of chemical shift perturbation changes upon binding of the three peptides suggested an overall conserved binding interface. We focused on the residues that exhibited chemical shift changes of at least one standard deviation above average for all three peptides. The data suggested significant rearrangements of residues in the hydrophobic core of Nsp9 in the β1-β5 region while β6 and β7 remained unperturbed **(Figure 4E, Figure S6H-I)**. Importantly, chemical shifts for the C-terminal alpha helix were not observed as discussed previously^44,48^, leaving open the possibility that the peptides could interact with the helix without showing any perturbations in the NMR HSQC spectra. To determine whether the Nsp9 is a dimer in solution, we measured the *T*_1_ and *T*_2_ relaxation times, which report on global motion of individual residues of the protein (**Table S6**). The ratio of the two relaxation times, gives the rotational correlation time τ_c_, which was 15.3 ns for the free Nsp9 protein and 9.9 ns for the protein in complex with NOTCH4_1605-1620_ peptide corresponding to estimated molecular weights of 29 and 18.8 kDa respectively. Because the molecular weight calculation from τ_c_ is highly dependent on the shape of the molecule, the experiments provide an estimate of the size, which nonetheless correlates well with the predicted molecular weight for a Nsp9 homodimer (25.7 kDa) and a Nsp9 monomer bound to the NOTCH4 peptide (14.7 kDa) implying that the addition of the peptide disrupts formation of the Nsp9 homodimer. To further test whether the peptides bind via the C-terminal helix we constructed a Nsp9Δα construct, which is missing 20 residues of the C-terminus **(Table S1**). As expected, the Nsp9Δα did not interact with the FITC-NKRF_8-23_ probe **(Figure S1, Table S4)**, corroborating our hypothesis that the identified peptides bind to the C-terminal helix of Nsp9 and interfere with Nsp9 homodimer formation.

In summary, we identified short peptide binders that interact with Nsp9 via a GΦxΦ[GDP] motif, which also encompasses the GxxxG motif found in the C-terminal protein-protein interaction region of Nsp9. This motif is present in many human proteins (**Figure 2B, Figure 4B**), and the interaction with these proteins could affect viral replication. Using NMR, we located the peptide binding interface and showed that peptide binding perturbs packing of the Nsp9 hydrophobic core, consequently interfering with dimer formation. Since Nsp9 interacts with NiRAN through the same C-terminal helix, this interaction, which is critical for viral replication, should also be disrupted^22^.

### Identified peptide ligands directly interfere with the Nsp10-Nsp16 complex

Since the SARS-CoV-2 virus does not have access to the cellular mRNA capping machinery, its genome encodes a full set of enzymes that facilitate complete 5’ capping of viral RNA, thereby promoting translation and evasion of the immune response^22,49,50^. In the final step of this process the methyltransferase Nsp16 catalyzes the C2’-O methylation of the first nucleotide of the RNA, forming a fully functional ^7Me^GpppA_2′-O-Me_-RNA^22,51,52^. To catalyse this reaction Nsp16 forms a complex with Nsp10, which in turn activates Nsp16 by stabilizing the vital S-adenosylmethionine binding pocket^49,51^.

While ProP-PD selections against Nsp10 did not result in specific enrichment of peptides, we identified peptides from six proteins that bind to Nsp16 (**Table S2, Figure S2**) of which we selected three peptides for affinity measurements. These peptides were derived from Dual specificity tyrosine-phosphorylation-regulated kinase 1B (DYRK1B_395-410_; PGHSPADYLRFQDLVL), Islet cell autoantigen 1-like protein (ICA1L_450-465_; NQDMSAWFNLFADLDP), and uncharacterized protein FLJ43738 (FLJ43738_76-91_; EDPLDSYLNFQALISP). Using the FP assay, we confirmed the interactions and determined that the peptides interacted with Nsp16 with *K*_D_ values of 56 μM, 30 μM and 62 μM, respectively (**Figure 5A**). Interestingly, the tightest binder, FITC-labelled DYRK1B (FITC-DYRK1B_394-409_), bound with a *K*_D_ of ~3 μM, exhibiting a 10-fold higher affinity compared to the unlabeled peptide with the same sequence (DYRK1B_394-409_) **(Figure S1, Table S4)** implying that the FITC label itself contributes to binding. Alanine scan SPOT array analysis of ICA1L_450-465_ and DYRK1B_395-410_ peptides revealed the presence of a hydrophobic motif (WxxxF) in the ICA1L_450-465_ peptide, which is also present in the peptide from the homologous ICA1 (**Figure 5B, Figure S2**). For the DYRK1B_395-410_ peptide the SPOT array analysis established the importance of a phenylalanine in P1 but did not confirm the second position of the hydrophobic motif (P5, which in this case would be a valine (**Figure 5B**, **Figure S2**). A putative [WF]xxxΦ motif is also present in the validated binding peptide from FLJ43738_76-91_ (Figure S2). ColabFold predicted that the three validated peptides form an alpha helix and bind to the hydrophobic groove formed by alpha helices α3, α4 and α10 of Nsp16 (**Figure 5C, Figure S7**). This result correlated with the SPOT array data and corroborated the importance of hydrophobic residues at the P1 and P5 positions of the Nsp16 binding motif.

**Figure 5:**
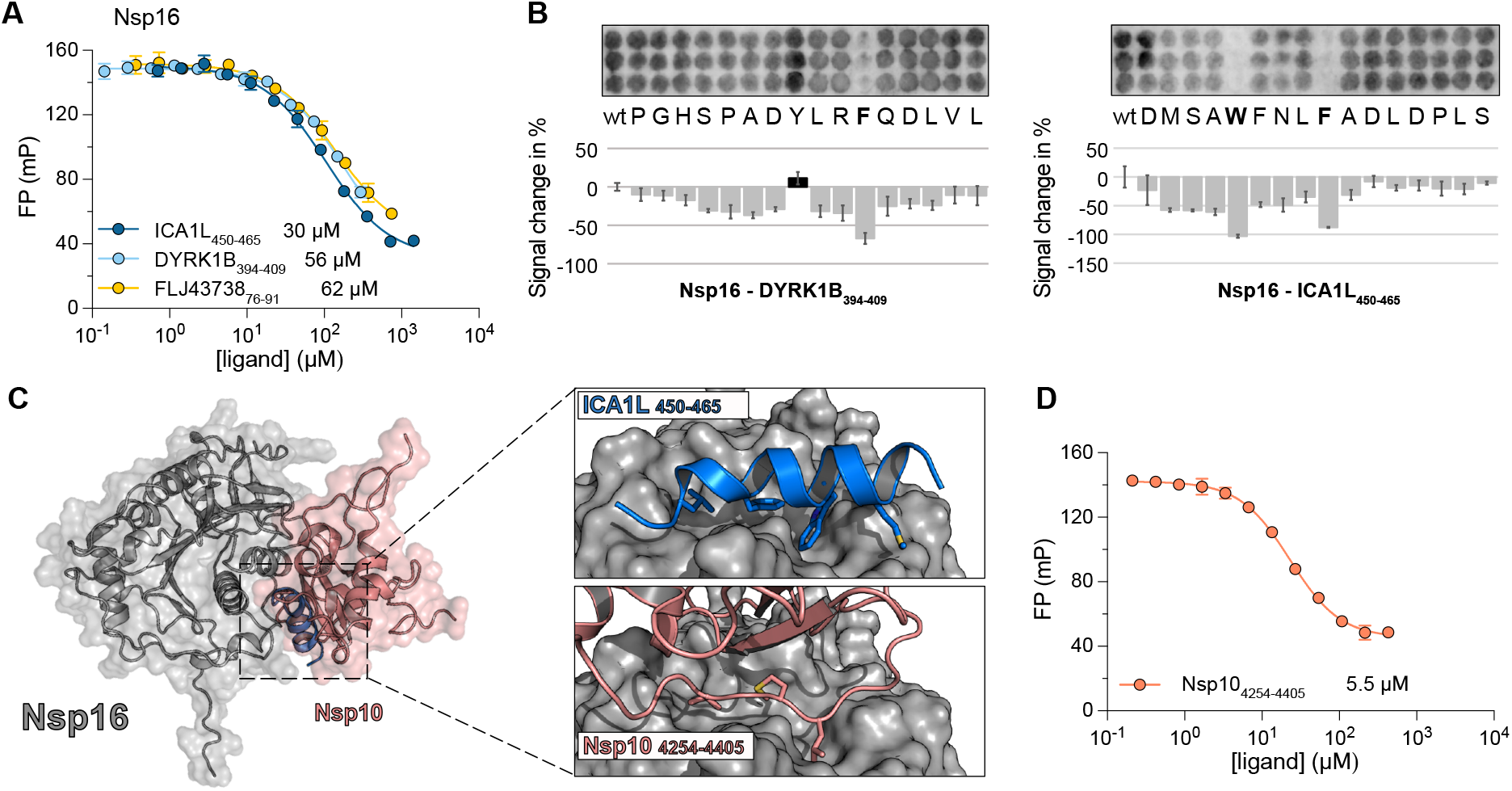
Peptides identified with ProP-PD directly compete with Nsp10 for binding to Nsp16. **(A)** Displacement experiments measuring affinity between Nsp16, and three peptide ligands derived from human proteins. *K*_D_ values are shown next to each peptide. **(B)** SPOT array alanine scans for the DYRK1B_395-410_ and ICA1L_450-465_ peptides. Residues involved in binding are shown in bold. Signal intensities were normalized to wild type (wt) and presented as percent signal change. **(C)** ColabFold structural predictions for the interaction between Nsp16 (grey), and the ICA1L_450-465_ peptide (blue). The predicted structure is superimposed onto the crystal structure of the Nsp16-Nsp10 complex (PDBid: 7jyy). The right panels show a magnified view of the peptide binding pocket highlighting the similarities of the Nsp16 binding to the peptide or Nsp10, respectively. **(D)** FP-monitored displacement by Nsp10 of a complex between Nsp16 and FITC-DYRK1B_394-409_ shows that Nsp10 and the peptide binds to the same surface of Nsp16. The *K*_D_ value for the Nsp10-Nsp16 interaction is indicated.

Since the same hydrophobic interface of Nsp16 facilitates the interaction with Nsp10^52^ we hypothesized that our newly characterized peptide ligands could compete with Nsp10 for binding to Nsp16 (**Figure 5C**). Indeed, we demonstrated that full-length Nsp10 directly competes with the labelled FITC-DYRK1B_394-409_ peptide in a displacement experiment (**Figure 5D**). This experiment also allowed us to determine the *K*_D_ of the interaction between Nsp10 and Nsp16 to be 5.5 μM.

Overall, we identified several peptide binders that interact with Nsp16, measured their affinity, and determined their binding interface with Nsp16. The peptide ligands compete with Nsp10 for binding to Nsp16 and could therefore act as inhibitors of the essential Nsp10-Nsp16 interaction, thereby hampering viral replication.

### Peptide ligands targeting Nsp3 ADRP, Nsp9 and Nsp16 inhibit viral proliferation

To investigate the role of the identified protein-protein interactions in viral replication, we selected 11 of the peptide ligands for further experiments in virus infection assays. We designed lentiviral expression constructs, expressing four repeats of each peptide interspaced by flexible GlySerThr linkers and conjugated C-terminally to an enhanced green fluorescent protein **(Table S7)**. VeroE6 cells were first transduced with the lentiviruses and, after 72 hours, infected with SARS-CoV-2 for 16 hours, after which the number of infected cells was measured. We found that five of the lentiviral constructs inhibited the production of viral particles, namely the Nsp3 ADRP binding EGFP-PARP10 construct, the Nsp9 binding EGFP-NOTCH4 and EGFP-LMTK3 constructs, and the Nsp16 binding EGFP-DYRK1B and EGFP-ICA1L constructs (**Figure 6A-B)**. Additionally, we confirmed the interference of the ligands with viral replication by treating the infected cells with the NOTCH4 and DYRK1B-derived peptides fused to the cell-penetrating HIV Tat-derived sequence (YGRKKRRQRRRGSG) **(Table S8)**. These experiments confirmed the inhibitory effects of the Nsp9 binding NOTCH4, as well as the Nsp16 binding DYRK1B peptides (**Figure 6C**), establishing these ligands as potential starting points for the development of peptidomimetic inhibitors of SARS-CoV-2 infection. To investigate if the peptides specifically inhibit SARS-CoV-2 replication, we also treated human coronavirus 229E (HCoV-229E) infected MRC5 cells with the cell penetrating constructs. Interestingly, the peptides failed to inhibit HCoV-229E infection (**Figure S8**), suggesting a beta-coronavirus specific inhibitory effect.

**Figure 6.**
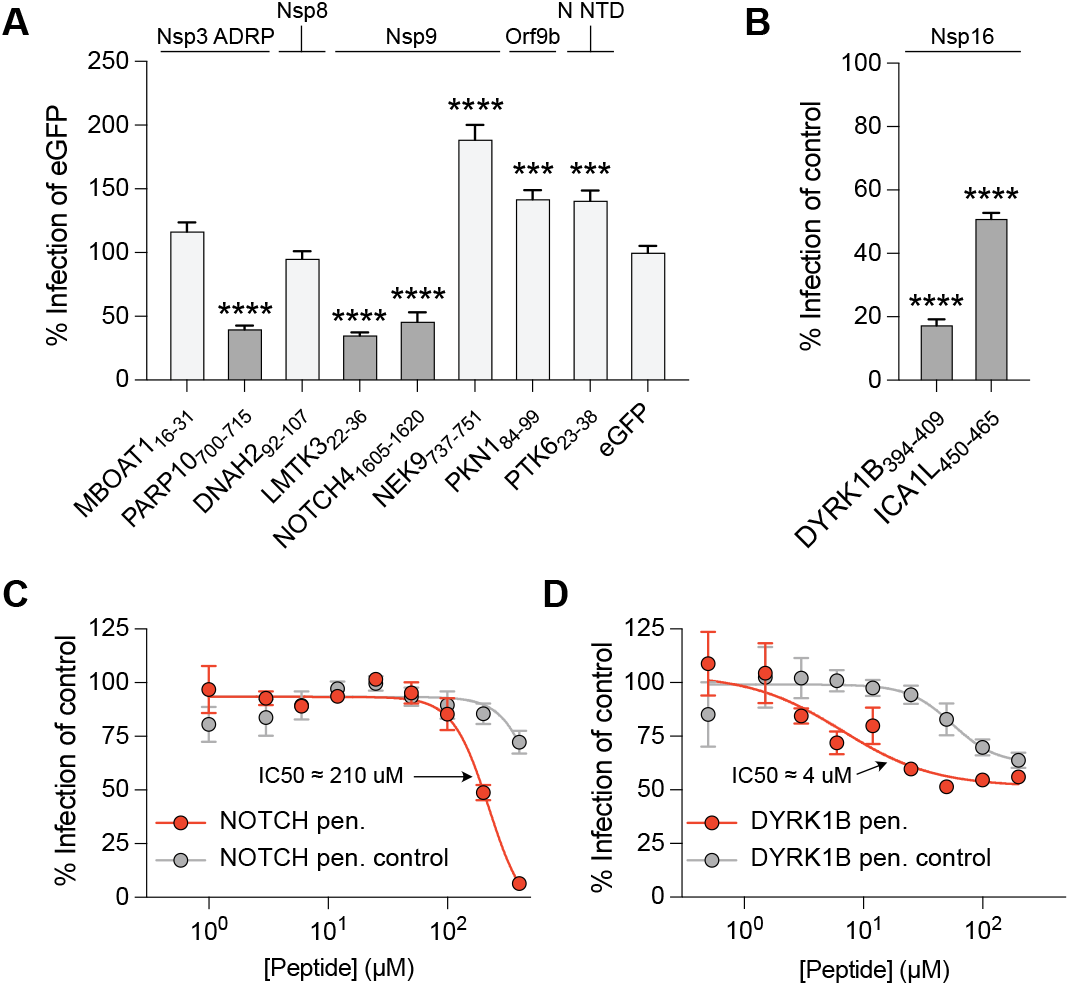
Inhibition of SARS-CoV-2 viral infection propagation by lentiviral constructs or cell penetrating peptides. **(A)** Inhibition is shown as percent infection of eGFP. Significance was determined using an unpaired t-test; *p< 0.05, **p< 0.01, ***p<0.001, and ****p<0.0001. Highlighted columns represent constructs, that showed significant inhibition. The target proteins of SARS-CoV-2 are stated above. **(B)** Inhibitions by lentiviral constructs expressing Nsp16-targeting peptides are shown as percent infection of control. In the control construct the key interacting residues are mutated to alanine. A complete list of the lentiviral constructs can be found in **Table S7**. Significance was determined using an unpaired t-test; *p< 0.05, **p< 0.01, ***p<0.001, and ****p<0.0001. **(C)** A serial dilution of either the cell penetrating NOTCH4 peptide (red dots) or the cell penetrating NOTCH4 control peptide (gray dots) was added to SARS-CoV-2 infected VeroE6 cells. The IC50 for the cell penetrating NOTCH4 peptide is indicated. **(D)** The same experiment was repeated but with either the cell penetrating DYRK1B peptide (red dots) or the cell penetrating DYRK1B control peptide (gray dots). The IC50 values shown were calculated using a sigmoidal dose response fitting equation.

## Discussion

In the present work, we performed a large-scale exploration of SLiM-mediated interactions between globular domains of SARS-CoV-2 proteins and peptides found in the intrinsically disordered regions of the human proteome. We aimed to answer three main questions: (1) How widespread are the SLiM-based interactions of viral proteins; (2) What are the human SLiM-containing binding partners of SARS-CoV-2 proteins; and (3) Can we exploit the newly identified peptide binders as antiviral agents.

Out of 26 SARS-CoV-2 protein domain constructs that we were able to express and purify, 11 showed phage enrichment from ProP-PD screens, and in total we identified 281 high/medium confidence human proteome-derived peptides that interact with SARS-CoV-2 protein domains. Several of the interactions between SARS-CoV-2 proteins are mediated through SLiMs and facilitate correct localization (Nsp3-N)^9^, complex formation (Nsp9-Nsp12)^22^, or regulate the activity of other viral proteins (Nsp10-Nsp16)^53,54^. Our results show that the same viral protein domains, which facilitate SLiM-mediated interactions between viral proteins, can also engage in interactions with human SLiMs. This may result in competition between host and viral SLiMs for binding to Nsp3, Nsp9 and Nsp16, potentially impacting the rate of infection processes such as viral RNA capping.

We found that Nsp9 can bind to several human proteins that contain GΦxΦ[GDP] motifs, many of which are associated with transcriptional regulation, suggesting a possible biological function of these interactions during the viral life cycle. While these findings are intriguing, it is not clear which human globular domain(s), if any, bind to the native, human GΦxΦ[GDP] motifs. Similarly, the C-terminal domain of the Ebola VP30 protein has been shown to bind to PPxPxY containing peptides found in both viral (N) and human proteins^10,55^. As with the Nsp9 binding GΦxΦ[GDP] motif, it is not yet known which human protein(s) binds to the PPxPxY motif in a functional context. The discovery of such human SLiMs binding to viral proteins can provide an alternative starting point in the search for novel SLiM binding domains and thus contribute to our understanding of human SLiM based interactions.

Since the start of the pandemic, there has been a rush to characterize SARS-CoV-2 protein interactions. Many of the large-scale studies have employed low-resolution techniques such as mass spectroscopy^5,27,30,47,56–58^ that provide information on binary interactions as well as on larger complexes. Furthermore, several studies aimed to identify potential inhibitors of viral infection, mainly targeting RNA-dependent-RNA polymerase^59–62^ and the proteases^63–65^, or producing antibodies against spike protein^66–70^. However, to increase the likelihood of successful drug development, efforts need to be extended to include other SARS-CoV-2 proteins. The Nsp9 dimer interface in the C-terminal helix with the GxxxG motif at its core has been proposed as a valid target for inhibitor development^71,72^. Our results confirmed that the Nsp9 binding GxxxG-containing peptides from LMTK3 and NOTCH4 indeed have antiviral effects, confirming the validity of the approach. The coronavirus Nsp10/Nsp16 interaction has also been shown to be a valid target for antiviral peptides, using Nsp10-derived peptides^53,54,73^. Similarly, we found that our NSP16 binding peptides had an antiviral effect. Other strategies targeting the Nsp10/Nsp16 complex include methyl donor site^51,71^ and RNA binding site targeting, as reviewed recently^74^.

In summary, we have demonstrated that ProP-PD screening is a viable strategy for identification of human peptides that bind to the globular domains of the SARS-CoV-2 proteome. We also showed that a subset of identified ligands inhibited viral replication in cell culture, and that these peptides could be successfully converted into cell-penetrating anti-viral inhibitors. Thus, our study expands the available peptide repertoire that may be used as starting point for future drug discovery targeting coronaviruses.

## Experimental procedures

### Protein expression and purification

The cDNAs encoding SARS-CoV-2 protein domains were ordered from Genescript in pETM33 expression plasmids (Table S1). The plasmid encoded an N-terminal His-tagged GST, a PreScisson protease cleavage site and the SARS-CoV-2 protein of interest. The proteins were expressed in BL21(DE3) gold *E. coli*. Bacteria was grown in 2YT medium (16 mg/ml peptone, 10 mg/ml yeast extract, 5 mg/ml NaCl) supplemented with 50 μg/ml kanamycin until OD_600_ reached 0.6, when the protein expression was induced by the addition of 0.5 mM isopropyl β-D-1-thiogalactopyranoside (IPTG). Proteins were expressed overnight at 18°C, bacterial cultures were harvested by centrifugation (4,500 g, 10 minutes) at 4°C and resuspended in lysis buffer A (50 mM Tris/HCl pH 7.8, 300 mM NaCl, 10 μg/ml DNase I and RNase, 4 mM MgCl_2_, 2 mM CaCl_2_ and cOmplete EDTA-free Protease Inhibitor Cocktail). For ProP-PD selections, protein domains were purified on a GSH affinity column (Pierce glutathione agarose) according to the manufacturer’s instructions. Following elution with 10 mM GSH in buffer A, the sample was used in selections where the His-GST moiety was immobilized on the plate according to the protocol described previously^25^. For fluorescence polarization experiments, after the initial GSH affinity purification step, the His-GST was cleaved off by PreScission protease (1:100 dilution; produced in-house) overnight at 4 °C. Following the cleavage, the samples were applied on a nickel Sepharose excel resin column to remove the His-GST tag. The SARS-CoV-2 protein was recovered in unbound fraction from the nickel column. The purity of the samples was inspected by SDS-PAGE and if needed an additional purification step was introduced, where the SARS-CoV-2 protein sample was applied to a size-exclusion chromatography column (Superdex 75, Cytiva). The identity of pure protein samples was confirmed with matrix-assisted laser desorption/ionization time-of-flight mass spectrometry (MALDI-TOF MS). Finally, protein samples were dialyzed against 50 mM potassium phosphate buffer pH 7.4 and flash frozen until further use.

For NMR experiments the Nsp9 expression construct was expressed in M9 minimal medium containing 1 g/l ^15^NH_4_Cl. After OD_600_ reached 0.6 the protein expression was induced with 1 mM IPTG, and expressed overnight at 18 °C. After expression, the purification protocol was the same as described above.

### Proteomic peptide phage display

We recently published the design of a Pro-PD library expressing disordered regions found in the human proteome and a pipeline to quickly analyze data from deep sequencing of enriched phages^25,75^. Briefly, GST-tagged bait proteins were immobilized in a 96-well Flat bottom Nunc MaxiSorp plates (Nunc, Roskilde, Denmark) at 4°C for 18 hours (10 μg per protein in 100 μl PBS). GST was immobilized in adjacent well as control. After immobilization the wells were blocked with blocking buffer (0.5% (w/v) BSA in PBS) for 1 hour at 4°C and washed 4 times with PT buffer (PBS supplemented with 0.05% Tween20). The phage library (prepared according to Benz et. al.^25^) was added (100 μl in PBS supplemented with 1 mM DTT) to the wells with immobilized GST for 1 hour, to remove nonspecifically binding phages, before being transferred to the wells with bait proteins. After 2 hours the unbound phages were removed by washing the wells 5 times with 200 μl of PT buffer, and the bound phages were eluted by the addition of 100 μl of E. coli OmiMAX in log phase with subsequent incubation for 30 minutes at 37°C. Bacteria were hyperinfected by the addition of M13KO7 helper phage and further incubated for 45 minutes at 37 °C. After successful infection, 100 μl of bacteria were transferred into 1 ml of 2xYT media supplemented with 100 μg/ml carbenicillin, 30 μg/ml kanamycin and 0.3 mM IPTG, followed by incubation for 18 hours at 37 °C under agitation. Next, bacteria were pelleted at 2000 g for 10 minutes and phage supernatant was transferred to a fresh 96-well plate, where the pH was adjusted by the addition of 1/10 volume of 10 x PBS and 1 mM DTT (final concentration), and heat inactivated at 65°C for 10 minutes. The resulting phages were used next day as in-phages in the second round of selection, where the whole procedure was repeated. To ensure high enrichment of binding phages the selection procedure was repeated four times. After the final day of selection, the phage enrichment was evaluated by pooled phage ELISA in 384-well Flat bottom Nunc MaxiSorp plates (Nunc, Roskilde, Denmark). Bait proteins and GST control were immobilized (5 μg in 50 μl PBS per well) at 4°C for 18 hours followed by blocking of the remaining well surface with 100 μl of 0.5% BSA in PBS, at 4 °C for 1 hour. Enriched phages from third and fourth rounds of selections (50 μl) were incubated with the corresponding bait protein for 1hour and the unbound phages were washed away 4 times with 100 μl PT buffer. Bound phages were incubated for 1 hour with 50 μl of M13 HRP-conjugated M13 bacteriophage antibody (Sino Biological Inc; 1:5,000 diluted in PT, 0.05% BSA). The wells were again washed 4 times with PT buffer followed by one wash with PBS. Substrate was added (40 μl of TMB substrate, Seracare KPL) and the enzymatic reaction was quenched by the addition of 40 μl of 0.6 M sulfuric acid. Finally, the absorbance at 450 nm was measured to quantify the phage enrichment with an iD5 plate reader (Molecular Devices).

### NGS analysis and ProP-PD data analysis

Peptide-coding regions were PCR-amplified and barcoded from the binding-enriched phage pools (5 μL) using Phusion High-Fidelity polymerase (Thermo Scientific). PCR products were normalized using Mag-bind Total Pure NGS, purified from a 2% agarose gel (QIAquick Gel extraction Kit), and analyzed using Illumina MiSeq v3 (1×150 bp read setup, 20% PhiX). Reads were demultiplexed, adapter and barcode regions were trimmed, and sequences were translated into peptide sequences. Peptides were annotated using PepTools. Confidence levels were assigned based on four different criteria: occurrence in replicate selections, identification of overlapping peptide sequences, high counts, and occurrence of sequences matching consensus motifs, as previously outlined^25^.

Protein interaction networks were built with CytoScape 3.9.1^76^. Inkscape 1.2.1 (https://inkscape.org) and Python Matplotlib 3.5.1 library (https://matplotlib.org) were used to build the figures. For the comparison of ProP-PD results with other protein-protein interaction datasets the previously reported evidence of protein-protein interactions was obtained from Biogrid (release 4.4.212)^77^, VirHostNet (release 3.0)^78^, IMEx (release 1.4.11)^79^, IntAct (release 4.2.18)^80^ and MINT (August 2022)^81^

### Fluorescence polarization

Fluorescence polarization experiments were performed as previously described in detail^7^. Briefly, peptides were ordered either unlabeled or as FITC-labelled constructs from GeneCust. First, *K*_D_ of the FITC labelled peptide for a certain interaction was determined by varying the protein concentration at a constant FITC-peptide concentration and fitting a hyperbolic function to the data. Next, a displacement experiment was performed to determine *K*_D_ for the unlabelled peptide. Labelled peptide (10 nM) was pre-incubated with protein such that approximately 50-60% of the labelled peptide was bound to protein (Nsp3 Ubl1: 150 μM, Nsp3 ADRP: 75 μM, Nsp3 SUD-M: 100 μM, Nsp9: 17.4-25 μM, and Nsp16: 7 μM). Unlabelled peptide was then added at increasing concentration to compete out the binding of the labelled peptide. Results were analyzed in GrafPad Prism (version 9.4.1). From this data set, the sigmoidal dose-response fitting function was used to obtain IC50 values. These IC50 values were further converted to *K*_i_ (=*K*_D_) of unlabelled peptide as described by Nikolovska-Coleska et. al.^82^

### Nuclear magnetic resonance spectroscopy

Before NMR experiments the ^15^N Nsp9 sample was dialyzed into 50 mM potasium phosphate buffer pH 6,7 supplemented with and supplemented with 1mM TCEP and 10% D_2_O. The final concentration of the sample was between 175-225 μM. All NMR experiments were performed on a 600 MHz Avance Neo HD NMR spectrometer (Bruker) equipped with a 5 mm TCI cryogenic probe. All ^1^H-^15^N TROSY HSQC spectra were recorded at 25°C making use of the BEST pulse sequence with 512 points in the direct and 256 points in the indirect dimension. Two or four scans per datapoint were taken with the relaxation delay of 200 μs. Similar experiments were performed upon addition of the Nsp9 binding peptide ligands. Final concentrations of NKRF_8-23_, LMTK3_22-36_, and NOTCH4_1605-1620_ were 416 μM, 230 μM and 323 μM, respectively. All Spectra were processed with TopSpin 3.2 and subsequently analyzed by MestReNova 14.1.0. The chemical peak shifts of Nsp9 residues were assigned based on comparison with previous assignments^44,48^. The perturbation of the chemical peak shifts upon addition of the peptides were calculated using equation 1:

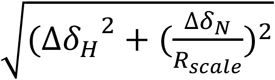

Where Δδ_H_ is chemical shift change in the hydrogen chemical shift dimension and Δδ_N_ is the chemical peak shift change in the nitrogen chemical shift dimension upon addition of peptide. R_scale_ is a scaling factor set to 6.5 as described before^83^.

To determine T1 and T2 relaxation times, TROSY HSQC based experiments were employed^84,85^ (PMID: 10729271 Zhu, PMID: 22689066). The overall rotational correlation time τ_c_ was estimated from the ratio of T1/T2 times (**Table S6**). The same Nsp9 - NOTCH4_1605-1620_ sample was used as for the previous experiments. For Nsp9 T1/T2 measurements fresh sample of Nsp9 was used at the concentration of 221 μM. The molecular weight of the species present in the sample was further estimated according to the equation 2: 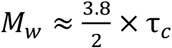 as described previously^86–88^.

### ColabFold predictions

ColabFold^34^ was used to predict binding of peptides to globular protein domains from SARS-CoV-2. ColabFold is based on AlphaFold2^89^ and AlphaFold2-multimer^90^. While the confidence scores varied for the predicted peptide conformations, all predictions for the structures of globular domains were in excellent agreement with solved crystal structure models (backbone alignment root mean square deviation between 0.4 and 1 Å).

### Alanine scanning SPOT arrays

20-mer, N-acetylated peptides on cellulose membranes were ordered from JPT (PepSpots). The membranes were activated with 5 ml methanol for 5 min at room temperature and washed with 10 ml TBST (50 mM Tris, 137 mM NaCl, 2.7 mM KCl, pH adjusted to 8.0 with HCl, 0.05% Tween-20) three times for 3 min at room temperature. The membranes were then incubated with 10 ml blocking buffer (5% skim milk powder (Merck Millipore, 115363) in TBST) for 2 hours at room temperature while rotated. The blocked membranes were incubated with concentrated GST-HA-tagged proteins of interest in blocking buffer overnight, at 4 °C, while rotated. After three quick rinses with ice-cold TBST, the membranes were incubated with HRP-conjugated anti-GST antibody (Cytiva, RPN1236) in blocking buffer for 1 hour at 4 °C, while rotated. Following three quick washes with ice-cold TBST, the chemiluminescent readout was carried out using ECL reagent (Clarity Max Western ECL substrate, 1705062, Bio-Rad) and ChemiDoc Imaging system (Bio-Rad). The acquired raw tiff images were analyzed using Image Studio Lite Ver. 5.2., and all values were normalized to the wild type results.

### SARS-CoV-2 infection assay

Vero E6 (*Cercopithecus aethiops*) cells (ATCC, CRL-1586) were cultured in Dulbecco’s modified Eagle’s medium (DMEM) (Sigma) supplemented with 10% fetal bovine serum (FBS) (HyClone) and 100 units/ml penicillin G with 100 μg/ml streptomycin solution (Gibco) at 37°C, 5% CO_2_, humidified chamber. MRC5, human lung fibroblast (ATCC CCL-171) cells were cultured in MEM medium (Gibco) supplemented with 10% FBS, and 100 units/ml penicillin G with 100 μg/ml streptomycin solution (Gibco) at 37 °C, 5 % CO_2_, humidified chamber. SARS CoV-2 (SARS-CoV-2/01/human2020/SWE accession no/GeneBank no MT093571.1, provided by the Public Health Agency of Sweden), was grown in VeroE6 cells, and used at passage number four. Human coronavirus 229E (HCoV-229E, ATCC CCL-171) was grown and titrated in MRC5 cells and used at passage one.

VeroE6 cells were transduced using the indicated lentiviruses as previously described^6^. After 72 h of transduction cells were infected with SARS CoV-2 for 16 h with MOI: 0.05 at 37 °C, 5% CO_2_. For peptide treatments cells were first infected with SARS CoV-2 MOI: 0.05 at 37 °C, 5% CO_2_, after 1 h of incubation the inoculum was replaced with medium containing the indicated concentration of peptide and cells were incubated at 37 °C, 5% CO_2_ for 16 h. After the infections, cells were fixed in 4% formaldehyde, permeabilized in 0.5% triton X-100 in PBS. Viral infected cells were revealed by staining using primary monoclonal antibodies directed against SARS CoV-2 nucleocapsid (SARS CoV-2, Sino Biological Inc., 40143-R001) or primary monoclonal mouse antibodies directed against J2 (HCoV-229E, Scicons 10010500), and secondary antibodies either donkey anti-mouse or donkey anti-rabbit IgG Alexa Fluor 555 secondary antibody (Invitrogen). Nuclei were counterstained with DAPI. Total cell number and number of infected cells/well were determined using a TROPHOS Plate RUNNER HD® (Dioscure, Marseille, France). Number of infected cells were normalized to DAPI count and presented as percentage infection of mutated control peptide for the lentivirus transduced cells or as percentage of mock treated cells in the case of peptide treatment. Results were analyzed in GraphPad Prism (version 9.4.1) and the sigmoidal dose-response fitting function was used to obtain IC50 values.

## Acknowledgements

This work was supported by the grants from the Swedish Foundation for Strategic research (Y.I., P.J.: SB16-0039), the Swedish Research Council (Y.I.: 2020-03380; PJ: 2020-04395; A.Ö.: 2018-05851), the Knut and Alice Wallenberg Foundation (Y.I., P.J., and A.Ö. via Science for Life Laboratory, KAW 2020.0241, V-2020-0699) and a Cancer Research UK Senior Cancer Research Fellowship (N.D.: C68484/A28159). We thank the medical faculty Umeå University strategic research resource and the Laboratory for Molecular Infection Medicine Sweden for generous support (A.Ö.), and the Biochemical Imaging Center at Umeå University and the National Microscopy Infrastructure, NMI (VR-RFI 2016-00968) for assistance in microscopy. Sequencing was performed by the SNP&SEQ Technology Platform in Stockholm. The facility is part of the National Genomic Infrastructure (NGI) Sweden and Science for Life Laboratory and is also supported by the Swedish Research Council and the Knut and Alice Wallenberg Foundation. We used the NMR Uppsala infrastructure, which is funded by the Department of Chemistry - BMC and the Disciplinary Domain of Medicine and Pharmacy, Uppsala University.

## Author contributions

FM, CB, ND, PJ and YI conceived the study. FM performed FP experiments. CB performed phage selections. EK performed SPOT arrays, RL performed viral assays, RI produced lentiviruses, HA, EA, CB and FM produced proteins, FM and CNC conducted NMR experiments and analyzed data. FM, LS, CB, ND, AKW, PJ and YI analyzed data. FM and PJ wrote the first draft. PJ and YI coordinated the study.

## Supplementary information

### Supplementary figures

**Figure S1.**
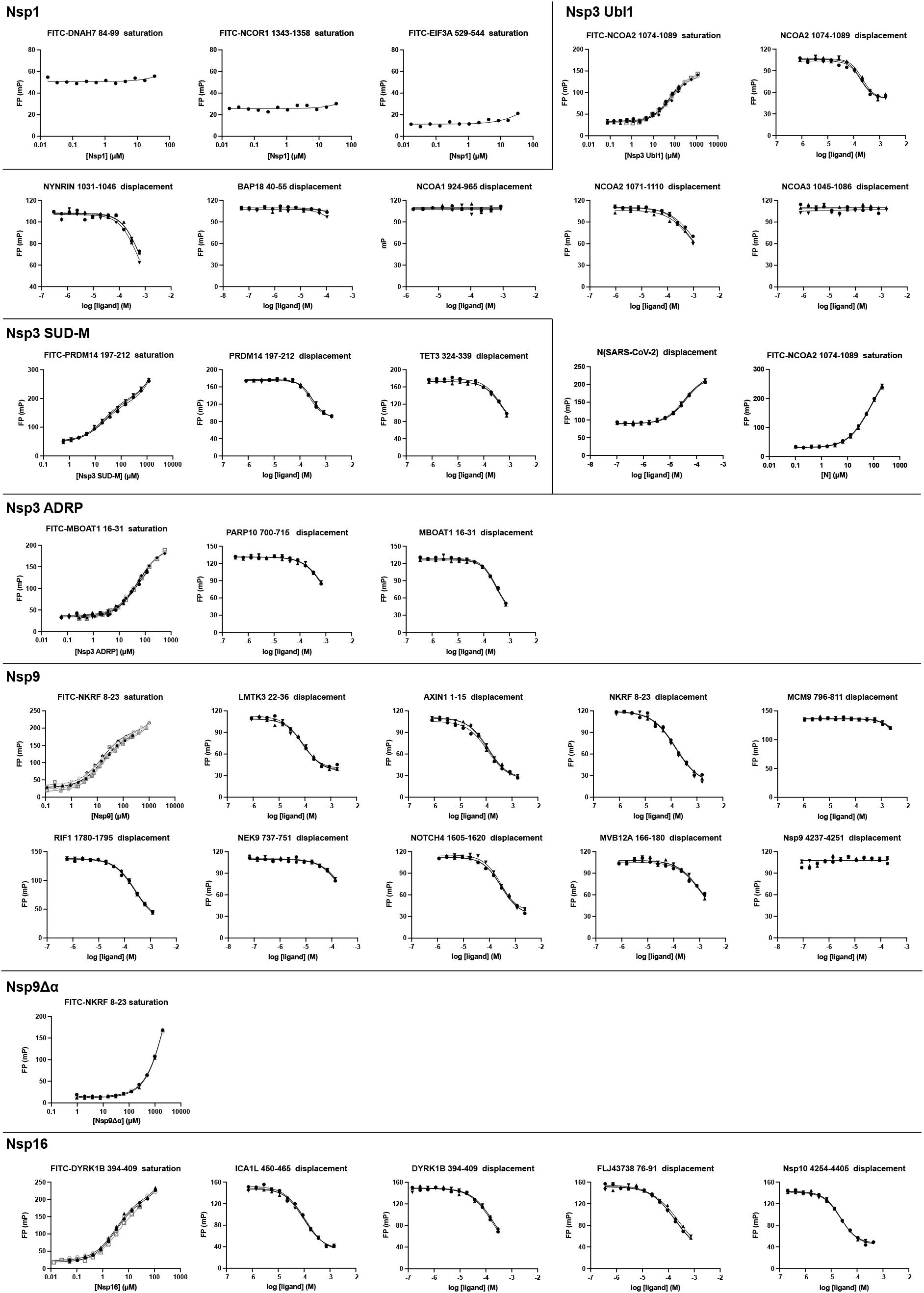
All affinity measurements performed in this study. Peptide sequences and calculated *K*_D_ values are presented in **Table S4**.

**Figure S2.**
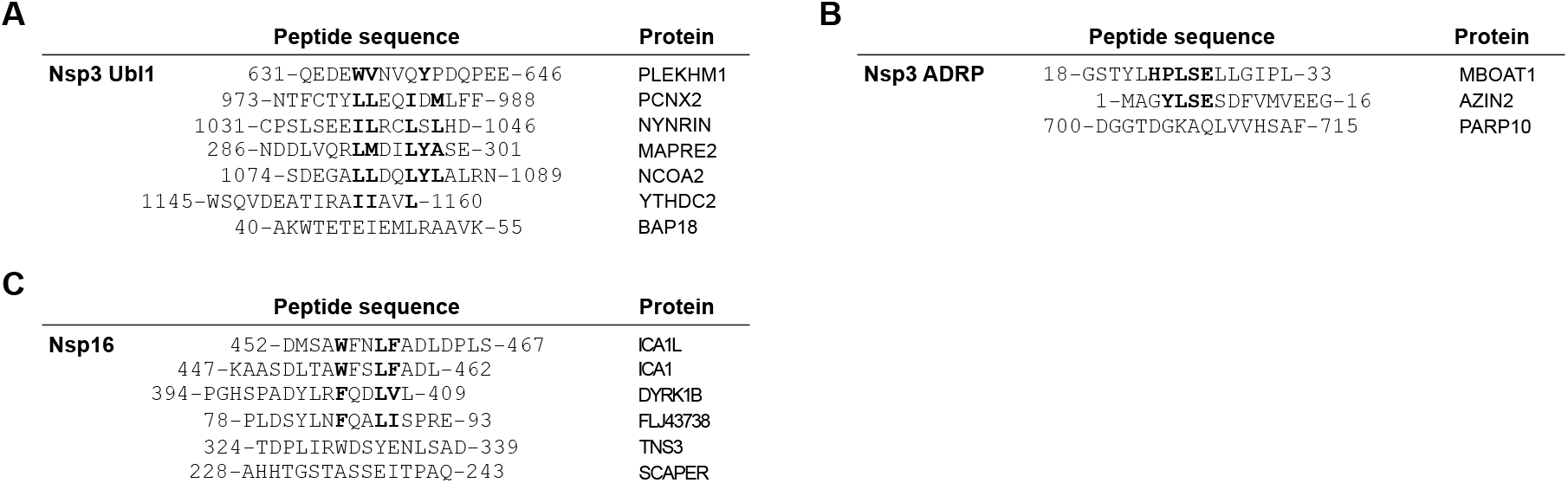
Alignment of the peptides identified by ProP-PD screens against. **(A)** Peptides identified by ProP-PD screens against Nsp3 Ubl1. Hydrophobic residues roughly corresponding to ΦxxΦxΦ are in bold. **(B)** Peptides identified by ProP-PD screens against Nsp3 ADRP. The common LSE motif is in bold. **(C)** Peptides identified by ProP-PD screens against Nsp16. The [W/F]xxxΦ motif is in bold.

**Figure S3.**
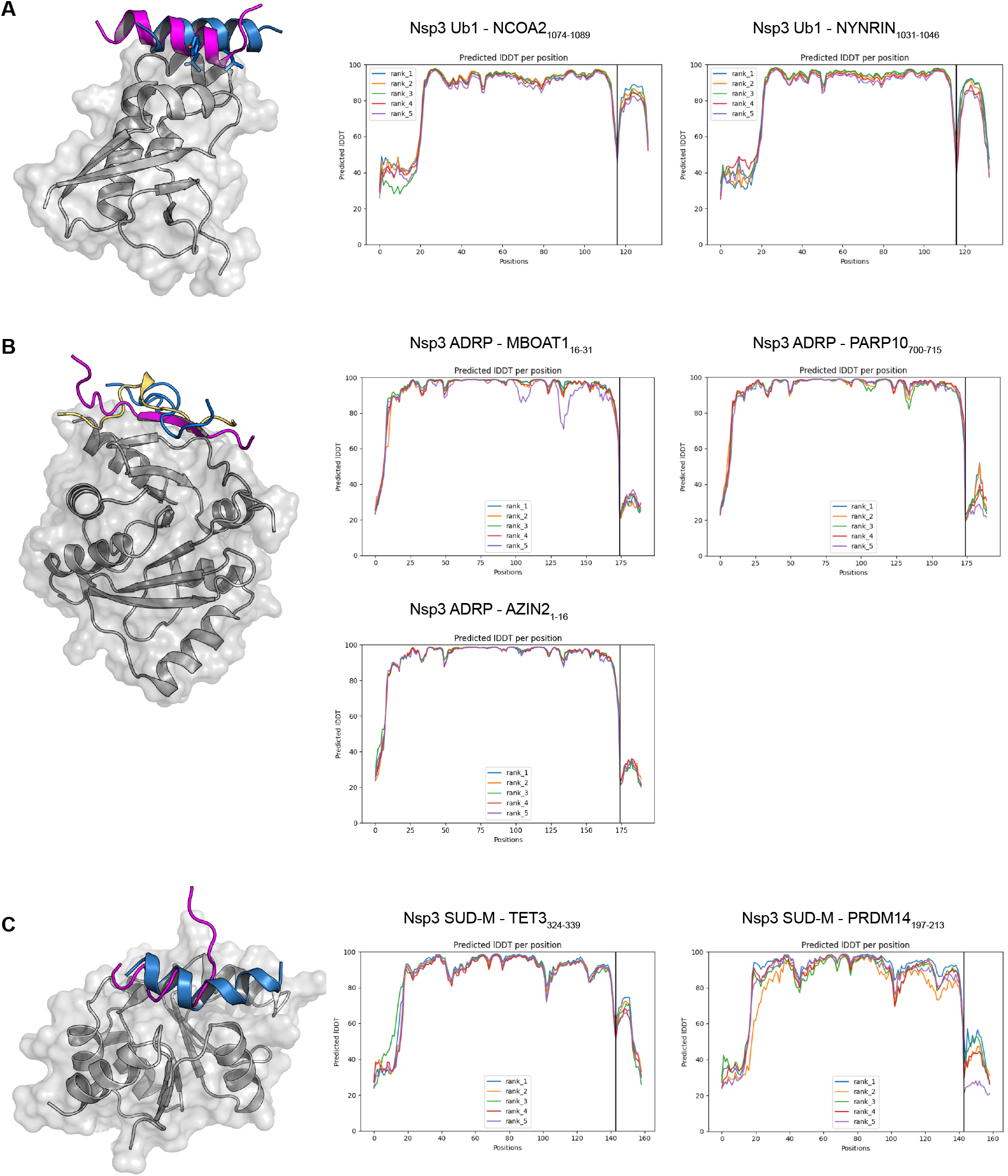
Models and confidence scores from ColabFold predictions for binding between Nsp3 globular domains and peptides identified in ProP-PD selections. **(A)** Model and confidence scores for prediction of Nsp3 Ubl1 in complex with peptides. The NCOA2_1074-1089_ peptide is in blue, and the NYNRIN_1031-1046_ peptide is purple. **(B)** Model and confidence scores for prediction of Nsp3 ADRP in complex with peptides. The MBOAT1_16-31_ peptide is in blue, the PARP10_700-715_ peptide is in purple, and the AZIN_1-16_ peptide is yellow. **(C)** Model and confidence scores for prediction of Nsp3 SUD-M in complex with peptides. The TET_324-339_ peptide is in blue, and the PRDM14_1074-1089_ peptide is purple.

**Figure S4.**
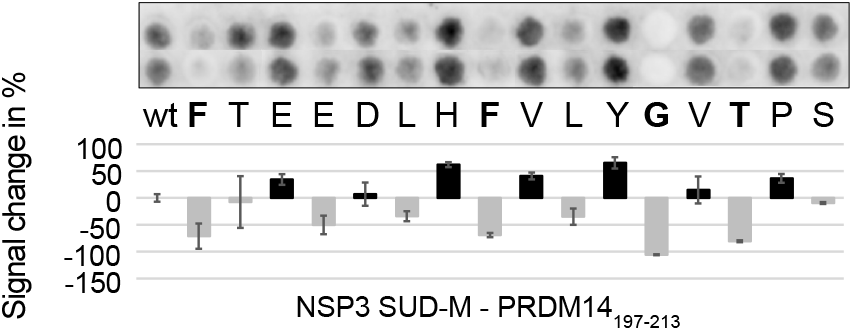
SPOT array alanine scans for the PRDM14_197-213_ peptide. The peptide was challenged with Nsp3 SUD-M. The residues where clear reduction of binding is observed are in bold.

**Figure S5.**
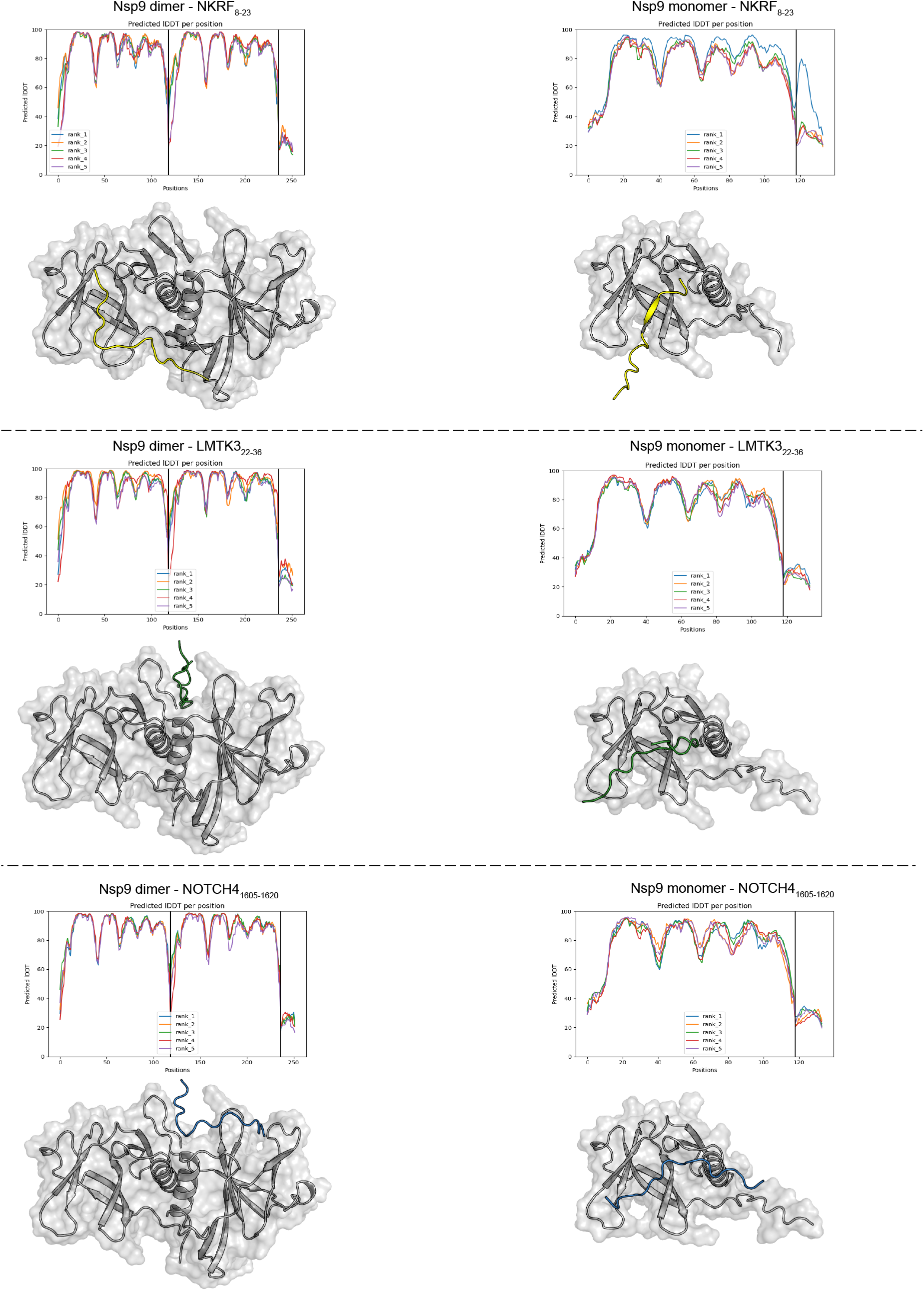
Models and confidence scores of ColabFold predictions for interaction between Nsp9 with NKRF_8-23_, LMTK3_22-36_, and NOTCH4_1605-1620_ peptides. For all three peptides the prediciton was run with either monomeric or dimeric Nsp9. The predicted peptides did not converge with either Nsp9 monomer or dimer model.

**Figure S6.**
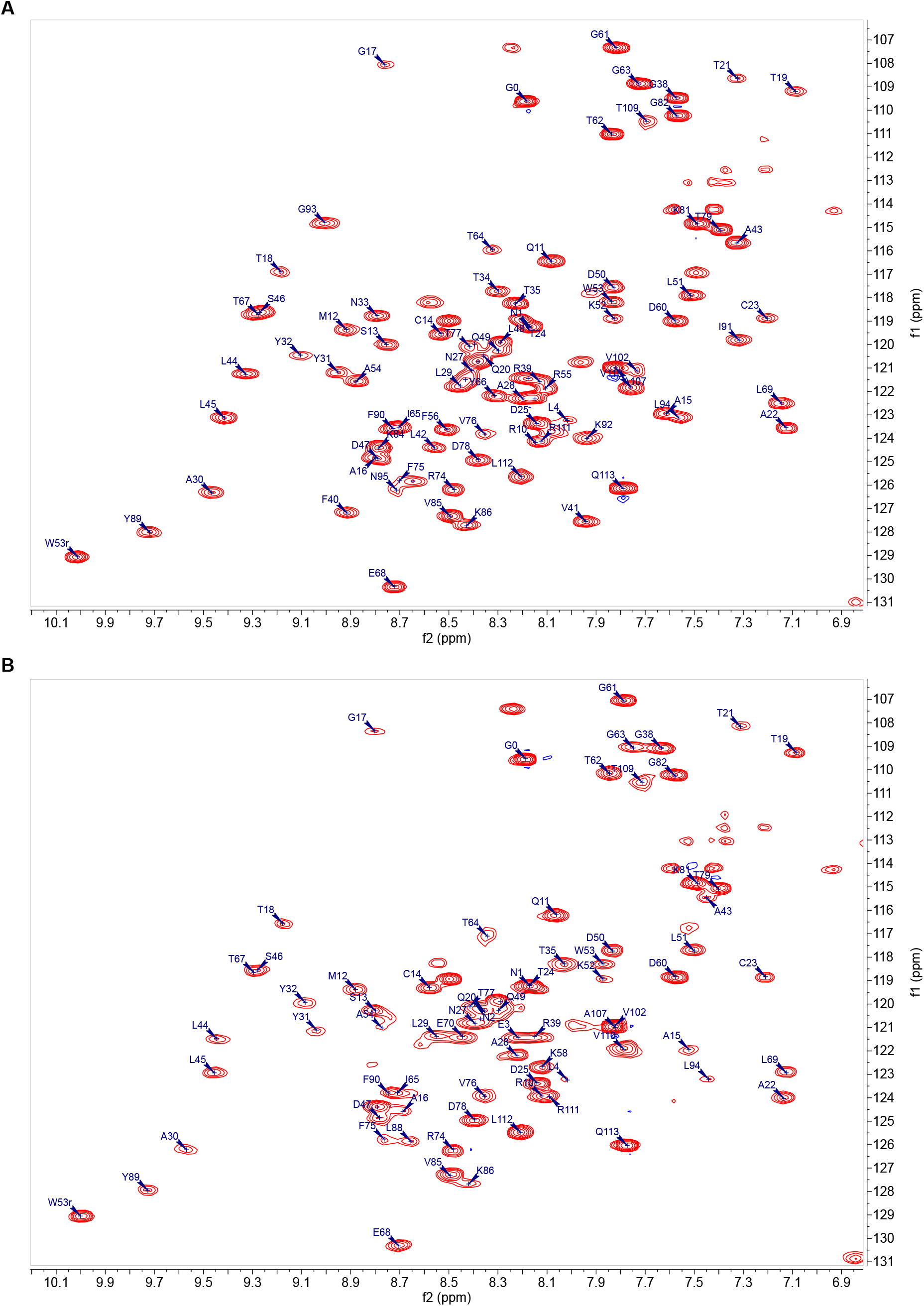
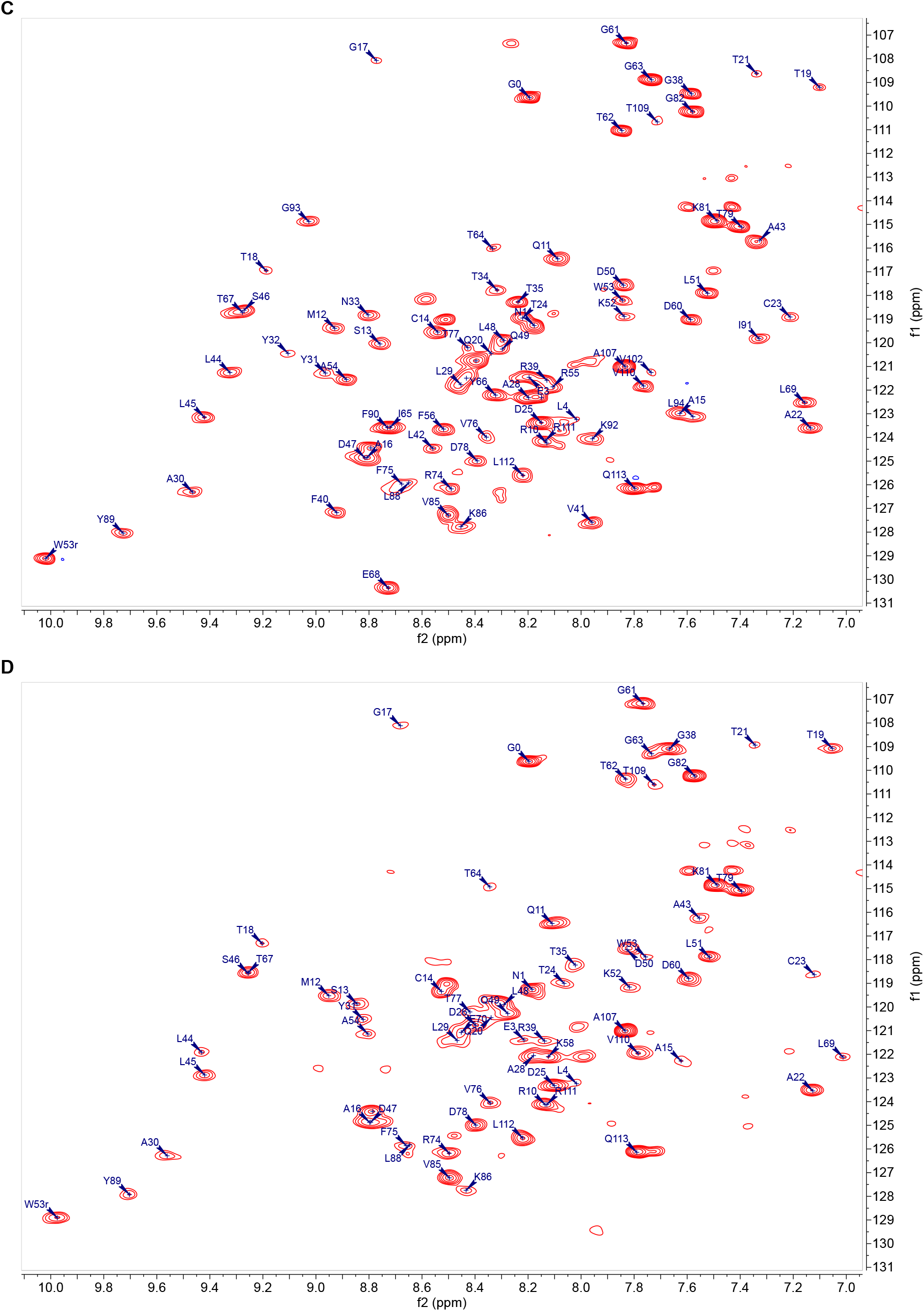
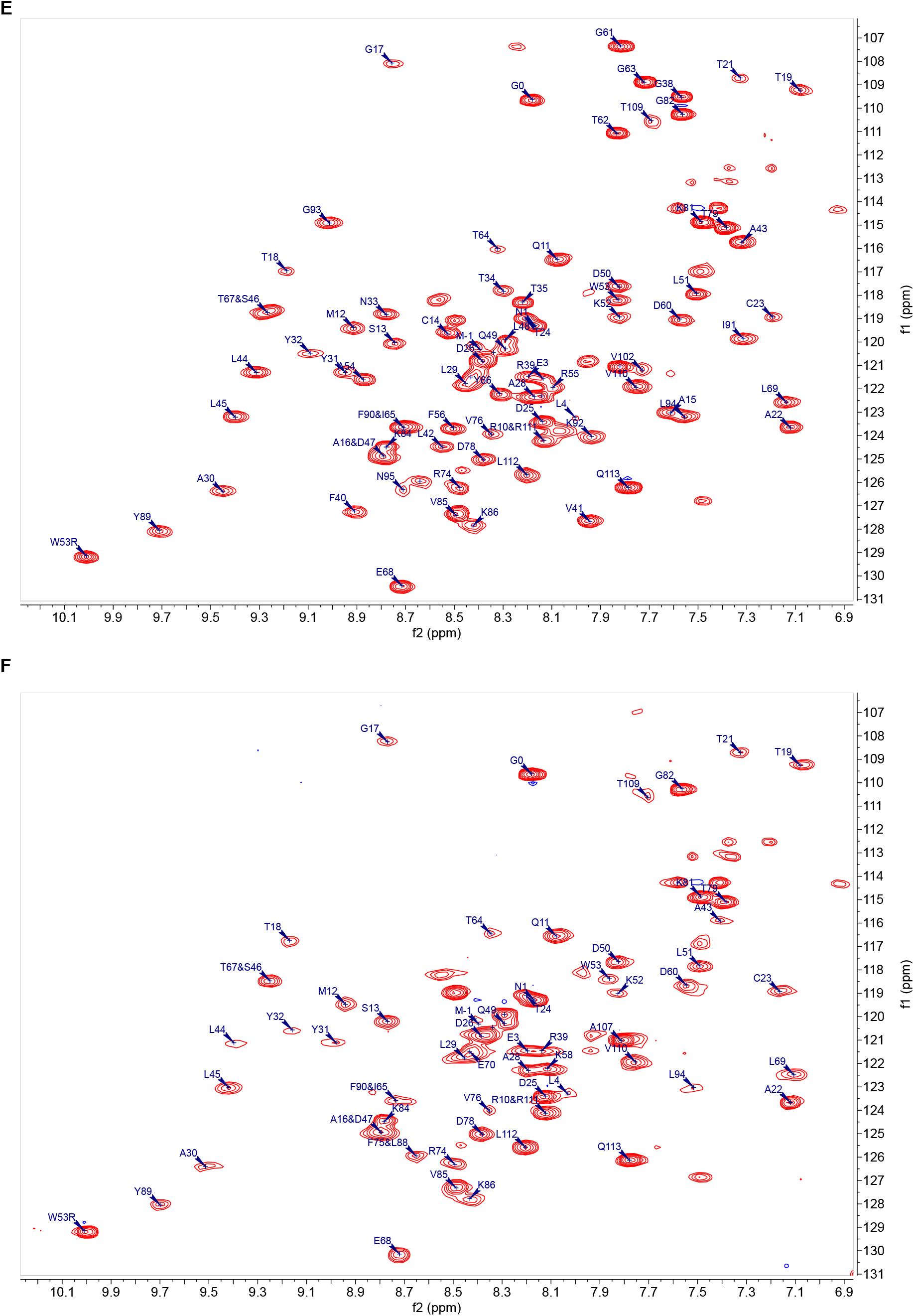
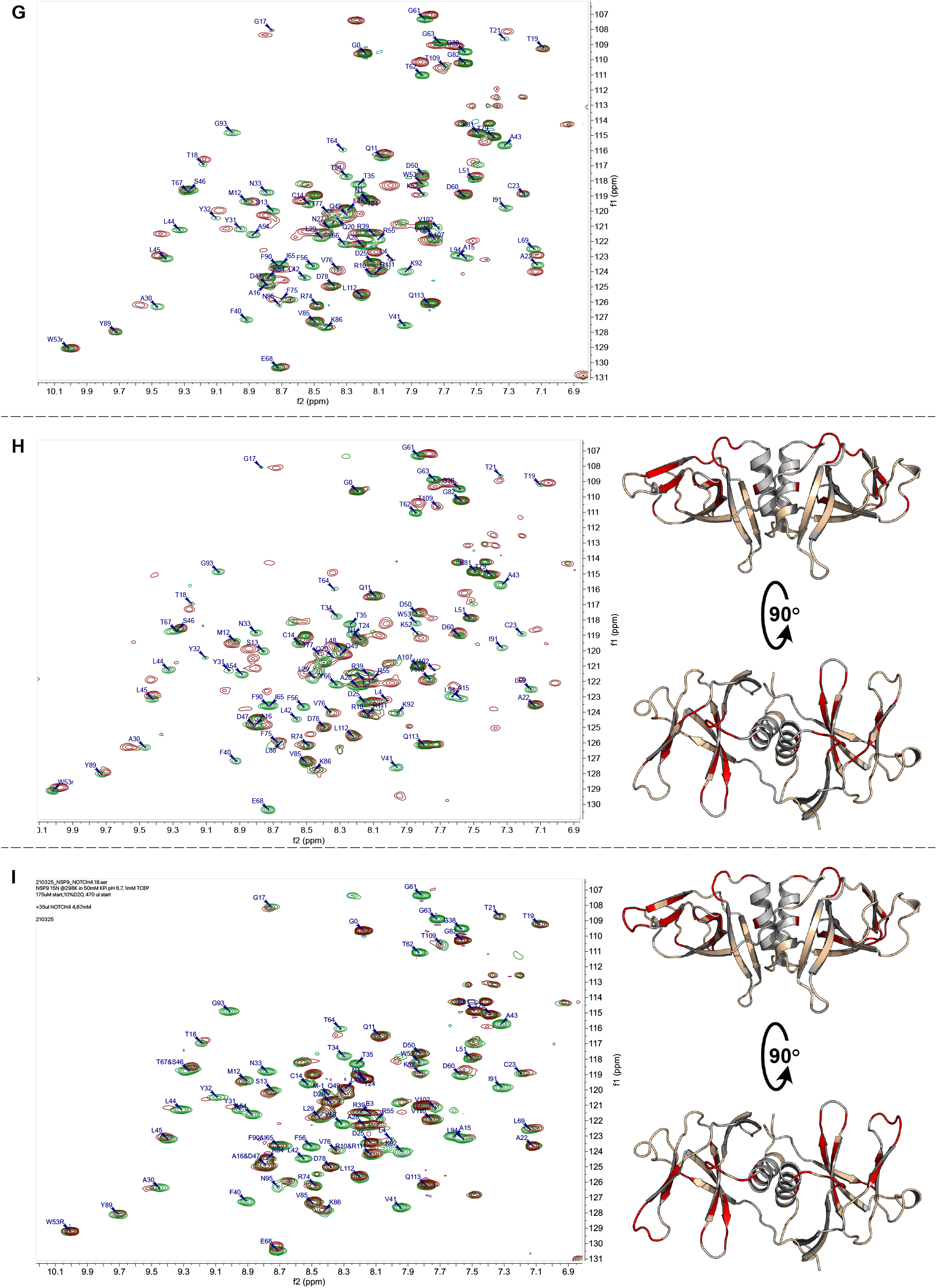
Raw HSQC spectra and Nsp9 model with highlighted peak shift changes for NKRF_8-23_, LMTK3_22-36_ and NOTCH4_1605-1620_ peptides. **(A)** Chemical shift annotation for Nsp9 without added peptide for the experiment where NKRF_8-23_ was added to the Nsp9. **(B)** Chemical shift annotation for Nsp9 with NKRF_8-23_ peptide. **(C)** Chemical shift annotation for Nsp9 without peptide for the experiment where LMTK3_22-36_ was added to the Nsp9. **(D)** Chemical shift annotation for Nsp9 with LMTK3_22-36_ peptide. **(E)** Chemical shift annotation for Nsp9 without peptide for the experiment where NOTCH4_1605-1620_ was added to the Nsp9. **(F)** Chemical shift annotation for Nsp9 with NOTCH4_1605-1620_ peptide. **(G)** Overlay of spectra from **(A)** and **(B)**. **(H)** Overlay of spectra from **(C)** and **(D)**. On the right side the residues of Nsp9 with a chemical shift change larger than one standard deviation upon addition of LMTK3_22-36_ are highlighted in red, the residues whose shift is bellow that threshold are in beige and the unassigned residues in gray. **(I)** Overlay of spectra from **(E)** and **(F)**. On the right side, residues of Nsp9 with a chemical shift change larger than one standard deviation upon addition of NOTCH4_1605-1620_ are highlighted in red, the residues whose shift is bellow that threshold are in beige and the unassigned residues in gray.

**Figure S7.**
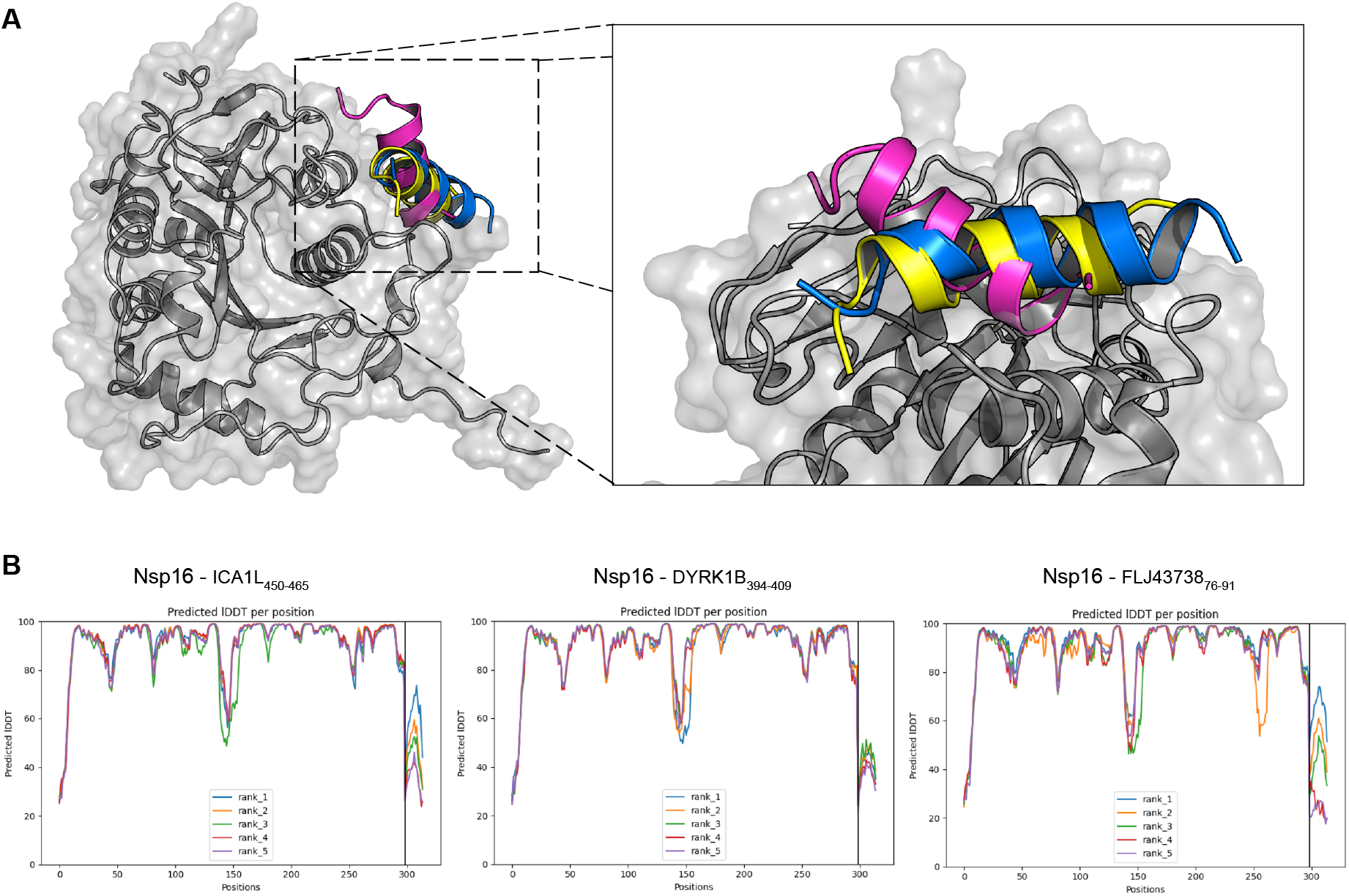
Models and confidence scores of ColabFold predictions for the interaction between Nsp16 and DYRK1B_395-410_, ICA1L_450-465_, and FLJ43738_76-91_ peptides. **(A)** Superimposed best predicted model for the interaction. The ICA1L_450-465_ peptide is in blue the DYRK1B_395-410_ peptide in purple and the FLJ43738_76-91_ peptide is in yellow. **(B)** Confidence scores for the predictions.

**Figure S8.**
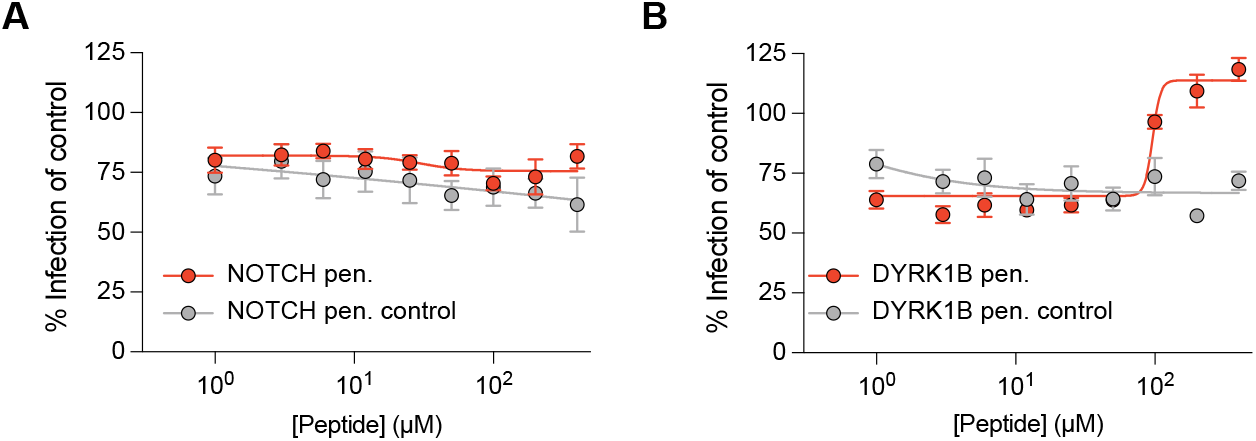
Cell penetrating peptides fail to inhibit HCoV 229E proliferation. **(A)** The MRC5 cells were infected with HCoV 229E and subsequently treated with either NOTCH4 cell penetrating peptide (red dots) or the NOTCH4 control cell penetrating peptide (gray dots). **(B)** The same experiments were performed with the exception that the cells were treated with either DYRK1B cell penetrating peptide (red dots) or DYRK1B control cell penetrating peptide (gray dots).

### Supplementary tables; see separate files

**Table S1. Overview of all SARS-CoV-2 protein domain constructs used in this study**.

**Table S2. ProP-PD selection results**

**Table S3. GO term analysis**

**Table S4. Overview of the peptides and affinity measurements performed in present work**.

**Table S5. NMR peak shift perturbation calculation**

**Table S6. NMR *T1, T2* calculation**

**Table S7. Lentiviral constructs used in this study**

**Table S8. Cell penetrating peptides**

